# Mutualist-Provided Resources Increase Susceptibility to Parasites

**DOI:** 10.1101/2023.10.15.562412

**Authors:** Eunnuri Yi, Nova Meng, Corlett W. Wood

**Affiliations:** Department of Biology, University of Pennsylvania, Philadelphia, PA, USA

**Keywords:** *Medicago truncatula*, mutualism, parasitism, indirect costs, symbiosis, nitrogen, resource use, legume, rhizobia, root-knot nematode, *Meloidogyne*, *Sinorhizobium*

## Abstract

**Background and Aims:**

- Mutualistic microbes often increase their host’s susceptibility to parasite infection, but the mechanism underlying this pattern remains unknown.
- We tested two competing hypotheses to identify the cause of this phenomenon.
- First, mutualist-provided resources could attract antagonists by making hosts more resource-rich.
- Second, mutualism establishment itself might increase host vulnerability to antagonists by enabling antagonists to exploit host defenses or immune systems, which are often downregulated during mutualism establishment.

**Methods:**

- To test which mechanism underlies increased parasite susceptibility, we experimentally decoupled mutualism establishment and mutualist-provided resources in the legume-rhizobia mutualism.
- We measured parasite load on *Medicago truncatula* plants infected with root-knot nematodes in a full-factorial design, in which we independently manipulated rhizobia inoculation (mutualism establishment) and nitrogen availability (mutualist-provided resources).

**Key Results:**

- Increased nitrogen availability increased parasite infection. However, its effect was non-linear and was not explained by nitrogen assimilation into plant tissues, indicating that this effect is not driven by parasite attraction to resource-rich hosts.
- We found little effect of mutualism establishment on susceptibility, as neither rhizobia inoculation nor nodulation without nitrogen fixation significantly increased parasite infection.

**Conclusions:**

- Our results suggest that mutualist-provided resources are an important driver of indirect ecological costs of mutualism, although the mechanism linking mutualist-provided resources and susceptibility to infection remains unknown.

## Introduction

Thousands of animal and plant species depend on microbes for crucial resources (Chaston and Goodrich-Blair 2010). Animal herbivores rely on gut microbes to digest recalcitrant plant tissue, and nearly 90% of land plants receive growth-limiting nutrients from fungi or bacteria (Smith 1991). However, these host-microbe mutualisms do not take place in a vacuum. Plants and animals host thousands of other microorganisms, many of which are parasites and pathogens (Christian *et al*. 2015). While many mutualisms protect their hosts by conferring resistance or enabling defense against antagonists, in some cases microbial mutualists make their hosts more susceptible to infection by parasites and pathogens (Toth *et al*. 1990; Miller 1993; Damiani *et al*. 2012; Salem *et al*. 2015; Wood *et al*. 2018). In this study, we capitalized on unique features of the mutualism between legumes and nitrogen-fixing bacteria to determine whether mutualist-provided resources or mutualism establishment increases the host’s susceptibility to infection.

Mutualistic species interactions are ubiquitous (Janzen and Boucher 1985). Mutualists provide fundamental biological functions for their partners and ecosystems (Bronstein 2001b; Traveset and Richardson 2014). These functions are diverse, spanning transportation, reproduction, nutrition, and protection against antagonists (Klepzig *et al*. 2009). For example, in the plant-mycorrhizae mutualism, mycorrhizal establishment primes plant immune responses, increasing resistance to herbivory and fungal disease (Jung *et al*. 2012; Yu *et al*. 2022). Fungal endophytes can also reduce herbivory on host grasses through the production of nitrogenous alkaloid compounds, although effects can vary in their benefit (Clay 1988; Faeth and Fagan 2002; Saikkonen *et al*. 2010). In the legume-rhizobia mutualism, *Crotalaria* legumes only synthesize pyrrolizidine alkaloid defenses in nodules formed by rhizobia; *Lotus* legumes inoculated with rhizobia have increased resistance to soil pathogens (Irmer *et al*. 2015; Prévitali *et al*. 2025). These three resource mutualisms provide additional defense benefits to their plant hosts.

However, mutualisms can also have adverse consequences for other species interactions. Mutualisms can mediate competition: plants with higher mycorrhizal colonization can suffer decreased biomass from increased competition (Schmitt and Holbrook 2003; Unger *et al*. 2021). Mutualisms can also compromise or conflict with other mutualisms: more effective ant bodyguards also deter host flower pollinators, leading to decreased seed set and individual mass (Ness 2006). Lastly, mutualists can attract antagonists like herbivores, predators, parasites, and pathogens (Adler and Bronstein 2004; Rice *et al*. 2021; Bell *et al*. 2022). Increasing susceptibility to antagonists is an especially well-documented indirect cost of mutualism, especially across multiple plant resource mutualisms involving mycorrhizae, *Serendipita* endophytes, and rhizobia bacteria (Hartley and Gange 2009; Opitz *et al*. 2024; Yin *et al*. 2025). Exploitation of mutualism by antagonists is widespread across taxa and ecosystems (Bronstein 2001a). The legume-rhizobia mutualism, ant-fungus nutritional mutualism, ant-plant protection mutualism, plant-pollinator mutualism and cleaner fish-client interactions are all exploited by pathogenic fungi, herbivores, parasites, and predators that imitate the host’s mutualist partner (Johnstone and Bshary 2002; Herrera *et al*. 2002; Frederickson and Gordon 2007; Little and Currie 2009; Meehan *et al*. 2009; Irwin *et al*. 2010; Wood *et al*. 2018).

There are two hypotheses to explain why mutualisms increase susceptibility to antagonists. First, resources provided by the mutualist may attract antagonists (Simonsen and Stinchcombe 2014). When key resources are difficult to obtain, antagonists may preferentially attack hosts with mutualists because these hosts are now a source of essential nutrients for the antagonist. For example, the fertilization of host plants by nitrogen-fixing rhizobia increases herbivory in *Medicago* legumes (Simonsen and Stinchcombe 2014). Endophyte mutualists increase root carbon which increased nematode parasite colonization (Opitz *et al*. 2024). Plants which invested more in colonization by mutualistic mycorrhizal fungi that provide phosphorus, nitrogen and other nutrients for their host (*Ambrosia*) supported higher insect performance (Yin *et al*. 2025). Similarly, a meta-analysis found that plants colonized by mycorrhizae were consumed more by chewing insects (Koricheva *et al*. 2009). This positive correlation between host investment in resource-provided mutualisms and antagonist presence has also been documented in the keystone marine mutualism between corals and zooxanthellae symbionts and corallivore antagonists (Rice *et al*. 2021).

Alternatively, the establishment of the mutualism may itself increase vulnerability to antagonists. We refer to this hypothesis as the mutualism establishment hypothesis. Mutualism establishment can open the door for antagonists to mimic mutualists or otherwise evade host defenses. The ant-plant mutualism is exploited by ant and beetle species that mimic ant mutualists and effectively function as parasites (Janzen 1975; Raine *et al*. 2004). A similar phenomenon has been documented in marine mutualisms, where the predatory fang blenny can mimic the cleaner wrasse to feed on tissues of fish visiting cleaning stations (Moland and Jones 2004). Many hosts also downregulate their immune systems when establishing mutualisms, increasing their susceptibility to infection (Salem *et al*. 2015). For example, *Arabidopsis* plants with *Serendipita* endophytes had altered systemic defense responses which increased cyst and root-knot nematode colonization (Opitz *et al*. 2024). Increased algal symbiont density was found to decrease the expression of immune-related transcripts in coral hosts (Fuess *et al*. 2020). Downregulated defense responses may be an especially important factor in symbiotic mutualisms, which involve the coordinated establishment of prolonged contact between partners (Mansfield and Gilmore 2019).

The above examples implicate both mutualist-provided resources and the establishment of mutualism in increasing the host’s susceptibility to parasites. A definite test of these competing hypotheses, however, requires de-confounding mutualism establishment from mutualist-provided resources in a single system. This is especially important because the two mechanisms are not mutually exclusive: both could (and probably do) operate at the same time. Nevertheless, the two mechanisms have very different implications for the evolution of mutualism. If resources provided by the mutualist increase susceptibility to antagonists, then selection imposed by antagonists may constrain the evolution of resource exchange in mutualisms. Alternatively, if establishment of the mutualism itself increases susceptibility to antagonists, then selection imposed by antagonists may favor the evolution of mechanisms to discriminate friend from foe, but would not affect the subsequent exchange of benefits.

The legume-rhizobia mutualism is ideal for testing these competing hypotheses because it allows for mutualist-provided resources (nitrogen) to be decoupled from establishment of the mutualism (inoculation leading to nodulation). The Mediterranean legume *Medicago truncatula,* with its rhizobial partner *Sinorhizobium meliloti,* is a model system for the study of resource exchange and symbiotic mutualisms (Cook 1999; Rose 2008). Bacteria from the Rhizobia family fix atmospheric nitrogen for leguminous plants in exchange for carbon (Shanmugam *et al*. 1978; Masson-Bolvin and Sachs 2018). To fix nitrogen, the rhizobia and plant must first form specialized structures called nodules through a tightly regulated process of signaling between the plant host and bacteria (Ferguson *et al*. 2019). Nodulation is a distinct stage of mutualism establishment that necessarily precedes resource exchange and is controlled by different plant and rhizobia genes (Fischer 1994; Lindström and Mousavi 2020). During nodulation, the bacteria enter the cell of the host root, dramatically changing their own development and the development of the host. Inside the nodule, rhizobia convert atmospheric nitrogen into ammonia, which is a bioavailable form of nitrogen for the legume host. *M. truncatula* associates with rhizobia in the genus *Sinorhizobium*, primarily *S. meliloti* and *S. medicae* (Harrison *et al*. 2017; Kearsley *et al*. 2024).

Furthermore, the rhizobia mutualism can increase the host plant’s susceptibility to the northern root-knot nematode *Meloidogyne hapla*, a generalist obligate endoparasite that establishes galls on many plant and crop species’ root systems (Castagnone-Sereno *et al*. 2013; Wood *et al*. 2018). Infective second-stage juvenile root-knot nematodes enter roots in the elongation region and migrate to their chosen spot, where they become sedentary and use their stylets to inject chemical effectors into host cells (Truong *et al*. 2015). These effectors modify host defense responses and cause plant cell nuclei to duplicate without cell division, creating multinucleated ‘giant cells’ which serve as metabolically active feeding sites and accumulate around the growing female to form galls (Bartlem *et al*. 2014). By feeding on carbon and nitrogen-rich phloem from these giant cells, root-knot nematodes act as strong nutrient sinks (Bartlem *et al*. 2014). Increased below-ground carbohydrate demand can lead to larger roots (e.g. Poll *et al*. 2007, Willig *et al*. 2023), while nematode damage to the nutrient transport function of the root vascular system can compromise absolute or relative shoot growth (e.g. Patil *et al*. 2012, Wang L et al. 2023). The resource extraction by root-knot nematode parasitism leads to a decrease in reproductive fitness in *M. truncatula* (Wood *et al*. 2018). A reproductive female nematode will lay an egg mass containing hundreds of eggs on the surface of each gall; these eggs hatch and reinfect the same plant host in a matter of weeks or months (Charchar and Santo 2009).

Both rhizobia-provided resources (nitrogen) and establishment of the rhizobia mutualism (inoculation leading to nodulation) could increase the susceptibility of *Medicago truncatula* to nematodes. Nitrogen is a commonly limited resource and an essential nutrient for various biological processes, and is known to increase attack by herbivores (Van Alstyne *et al*. 2009; Chen *et al*. 2010). Nodulation, however, dampens the plant’s defense response. Rhizobial Nod factors, essential for nodulation, bind to plant receptors and suppress plant immunity. One of these Nod factor receptors, previously thought to function solely during rhizobial symbiosis, is also active during plant pathogen infection, indicating that Nod receptors may play a broader role in discriminating between mutualists and antagonists (Gourion *et al*. 2015). Rhizobia and root-knot nematodes both form specialized root structures on *Medicago*, and the two symbionts have also been shown to indirectly interact through the legume host (Catella *et al*. 2022). In cases of co-colonization with rhizobia and nematodes, nematodes tend to reduce rhizobia nodule numbers, while rhizobia can have varying effects on nematode gall formation (Wood *et al*. 2018; Costa *et al*. 2021; Burr *et al*. 2022).

In this study, we independently manipulated mutualist-provided resources (nitrogen) and mutualism establishment (inoculation and nodulation) in the legume-rhizobia system to evaluate their relative impact on parasite infection. If mutualist-provided resources are the mechanism that affect parasite infection, we predict that nitrogen fertilization will increase parasite infection. Alternatively, if mutualist-provided resources are the mechanism that affect parasite infection, we predict that inoculation with rhizobia will increase parasite infection. In addition, we predict that plants that form more nodules will be more heavily infected by parasites, even if those nodules do not fix nitrogen. We found that fertilization with nitrogen affected parasite resistance, but nodulation with rhizobia did not. Our results suggest that mutualist-provided resources are an important driver of indirect ecological costs.

## Methods

### Greenhouse Experiment

We factorially manipulated rhizobia inoculation and nitrogen availability to disentangle the effects of mutualism establishment and mutualist-provided resources on susceptibility to parasite infection. Each treatment had between nine to 13 plant replicates. We manipulated mutualist-provided resources via the application of three different fertilizer formulations which primarily and dramatically varied in nitrogen (0, 0.625, and 5 mM N from nitrate, hereafter referred to as no-N, low-N, and high-N). Each fertilizer contained the same concentration of iron versenate, magnesium, phosphorus, and micronutrients (Mn, Cu, Zn, B, Mo). The recipes we used to prepare our fertilizers are commonly used in the legume-rhizobia literature to manipulate nitrogen (Table S9; Moreau *et al*. 2008; Batstone *et al*. 2017).

We manipulated mutualism establishment via two inoculation treatments: uninoculated with no rhizobia (R-; no nodulation) and inoculated with rhizobia (R+; nodulation). We used four different rhizobia strains (A145, Rm41, USDA1021, and WSM1022) in our experiment. Our selection of rhizobia strains was intended to maximize variation in nitrogen fixation (mutualist-provided resources) to deconfound nodulation and nitrogen fixation in inoculated plants. Rhizobia strains A145 and Rm41 can form small white nodules but are incompatible for nitrogen fixation with our chosen genotype, *M. truncatula* Jemalong A17; molecular and genetic analysis found that these strains lack the compatible *fix* gene (Wang *et al*. 2018). Rhizobia strains USDA1021 and WSM1022 both nodulate and fix nitrogen on *M. truncatula*.

We germinated seeds of *M. truncatula* genotype A17 (“Jemalong”) under sterile conditions in a sequence of scarification, sterilization, and stratification (Barker *et al*. 2006). We filled 66-mL “Cone-tainer”™ pots (Stuewe & Sons, Inc., Tangent, OR, USA) for planting with a 1:4 (volume:volume) mixture of perlite and sand. These filled “Cone-tainer” pots were sterilized by autoclaving at 121 C twice for 90 minutes. We planted one germinant in each “Cone-tainer” and fertilized each plant with 5 mL of Fahraeus fertilizer corresponding to its respective nitrogen treatment (Batstone *et al*. 2017). The experiment was set up in a temperature-controlled greenhouse at the University of Pennsylvania’s Carolyn Lynch Laboratory. We fertilized plants with 5 mL of fertilizer twice a week using a peristaltic pump with tubing that was sterilized before and after each use. We grew the plants in four complete blocks, where each block contained plants from every nitrogen treatment and every rhizobia treatment (Figure S6). We used 12-inch tall transparent plexiglass sheets to separate plants inoculated with different rhizobia strains to minimize the risk of cross-contamination.

Two weeks after planting, we inoculated all plants with nematodes and tryptone yeast media with or without rhizobia. To collect M. *hapla* nematode eggs, galled roots were collected from a cv. ‘Rutgers’ tomato plant infected with the LM line of *M. hapla* nematodes procured from North Carolina State University. This line was originally collected in France, the native range of *M. truncatula* (Guo *et al*. 2017). To extract eggs, we cut the roots, shook them in 10% bleach for two minutes, and strained them over stacked sieves with 75 and 25 μm pore size to filter large particles and collect eggs. After decanting the eggs off the bottom sieve with tap water, we counted samples of the collected eggs under a microscope to calculate egg density in solution. We then diluted the culture with water to inoculate each plant with approximately 300 nematode eggs suspended in 5 mL DI water.

To inoculate plants with rhizobia, we grew strains from glycerol stocks and stab cultures in sterile tryptone yeast liquid media in a shaking incubator at 180 rpm and 30 °C. After two days, we streaked the cultures onto solid tryptone yeast agar media in a biosafety cabinet. These plates were incubated at 30 °C until distinct colonies formed, at which point we selected individual colonies from each strain to grow in tryptone yeast liquid media. After these single-colony rhizobia cultures grew for two days in the shaking incubator, we diluted the liquid media to an OD600 of 0.1 using a spectrophotometer. Each plant in the rhizobia-positive R+ treatment was inoculated with 1 mL of diluted culture containing either A145, Rm41, USDA1021, or WSM1022. Plants in the R-treatment (rhizobia-free, uninoculated) received 1 mL of sterile tryptone yeast liquid media as a control.

### Data Collection

We harvested all plants 7.5 weeks after the start of the experiment (53 days; May 28-July 20, 2021). Shoots were dried in a drying oven at 60°C for at least a week before being weighed to the nearest 0.0001g (Mettler AE 100). We counted seed pods and flowers on dried shoots. Root tissue was stored in plastic bags at 4°C to preserve nodules and galls.

We counted nematode galls under a dissecting microscope to quantify nematode infection (Wood *et al*. 2018). A single reproductive female resides in a gall structure and will generally lay a single egg mass. However, multiple egg masses can be present on what looks to the human eye like a single continuous gall. We dyed root samples with red food coloring to highlight nematode egg masses, which we also counted (Kandouh *et al*. 2019). We found that galls and egg masses were extremely highly correlated, with each gall generally containing a single egg mass (r^2^=0.97; Figure S7). Galls were not removed before weighing roots, because it is prohibitively time-intensive to do so, and because it difficult to remove galls without removing adjacent root tissue. Although galls therefore contribute to total root weight, nematode galls are so small and constituted a very small fraction of total root area such that their effect on root size is almost certainly negligible relative to the effects of our experimental treatments.

We also counted rhizobia nodules under a dissecting microscope to quantify rhizobia colonization (Wood *et al*. 2018). We scored nodule color as a proxy for active nitrogen fixation, which is a common practice across the legume-rhizobia literature (Ott *et al*. 2005; Tajima et al. 2007; Crook et al. 2012; Wang et al. 2016; Forrester et al. 2020; Wendlandt et al. 2022). White nodules indicate the absence of active leghemoglobin, which is pink in color and is necessary for N-fixation by binding oxygen (Regus *et al*. 2017, Wang *et al*. 2016, 2019). We classified pink and yellow nodules as N-fixing, and white nodules as non-N-fixing. We did not observe any senesced nodules, which are typically dark brown. After classification, nodules were plucked off with dissecting tweezers and stored in 1mL microcentrifuge tubes at 4°C. Both roots and nodules were dried in a drying oven at 60 °C for at least a week and also weighed to the nearest 0.0001g.

Plants inoculated with rhizobia were generally well-nodulated, forming a median of 28 and an average of 34 nodules per plant (Figure S1A). Uninoculated plants formed a median of 0 nodules and a mean of 3.7 nodules. Of the 22 surviving plants in the uninoculated treatment, eight formed nodules; most of the rhizobia contamination was limited to a few nodules (1-7), suggesting that they were likely formed late in the experiment and contamination was minimal (Figure S1A). Only two uninoculated plants formed more than 13 nodules, the first quartile for rhizobia-inoculated plants. For all analyses involving rhizobia strain or comparison between inoculation statuses, we excluded the two uninoculated samples that formed 13 or more nodules. Contrary to our expectation, some plants inoculated with A145 and Rm41, the “non-N-fixing” strains, formed pink (nitrogen-fixing) nodules (Figure S1D; Table S1). It is possible that a small number of these N-fixing nodules on plants inoculated with A145 and Rm41 may have been due to contamination by other strains. However, the low nodule counts for the few contaminated uninoculated plants suggest that contamination would only account for a small proportion of these N-fixing nodules. It is unlikely that the lack of difference between Fix- and Fix+ strains was caused by significant cross-contamination between rhizobia treatments, because most uninoculated R-plants remained nodule-free and the four strains significantly differed in nodule number, nodule mass, and proportion of non-N-fixing and N-fixing strains (Figure S1, Table S1).

To quantify shoot nitrogen and carbon concentration, we performed elemental combustion analysis. Dried shoot tissue was pulverized using a mortar and pestle. Samples of homogenized shoot tissue were weighed on an ultra-micro balance (CP2P Microbalance, Sartorius GmbH, Goettingen, Germany) in 4-5 mg increments and packaged into tin capsules. For shoots smaller than 4 mg in weight, we packaged the entirety of the sample available, excluding samples that were smaller than 2 mg. Elemental combustion analysis was run on a Costech ECS4010 Elemental Analyzer (Costech Analytical Technologies, Inc. Valencia, CA, USA) using atropine as a standard. Each run consisted of about 45 shoot samples, with five atropine standards per run which were used to generate a C:N curve. With this curve, the elemental analyzer calculated carbon and nitrogen composition (%) for each sample. Thirty-six plant samples were replicated across runs, which were consistent (r^2^=0.999). By multiplying carbon and nitrogen composition by shoot biomass, we calculated the estimated total nitrogen in the shoot (Figure S2, Table S2).

### Statistical Analysis

We fit generalized linear mixed models in R version 4.5.1 and RStudio Version 2025.09.1+401 (2025.09.1+401) using the glmmTMB package (Brooks *et al*. 2017) assuming a normal (Gaussian) distribution unless otherwise noted. For models with nematode galls (our proxy for parasite load) as the response variable, we fit a zero-inflated negative binomial error distribution. Galls and egg masses were highly correlated (r^2^=0.97; Figure S7), and results were qualitatively similar when run with egg masses instead of galls (Figure S8, Table S10). We fit block as a fixed effect because our blocking design contained four levels (Figure S6), too few to accurately estimate the among-block variance (Gelman and Hill, 2007).

In preliminary analyses, we used rhizobia strain as a fixed effect, but found no significant variation across the four rhizobia strains in host benefit, which we assessed by measuring shoot biomass and shoot nitrogen concentration (Figure S2, Table S2). Strains differed in the number of nodules formed, total nodule mass, and proportions of N-fixing (yellow and pink) nodules (Figure S1, Table S1). Given that our designated Fix-strains (A145 and Rm41) did not measurably differ from our Fix+ (USDA1021 and WSM1022) strains in mutualist-provided nitrogen, we recognized that it would not be appropriate to compare non-N-fixing to N-fixing strains to test our initial hypothesis about the effect of mutualist-provided resources on parasite susceptibility. We thus performed two complementary analyses to test whether mutualist-provided resources (nitrogen) or mutualism establishment (inoculation leading to nodulation) increased parasite load.

First, we tested for an effect of our nitrogen and rhizobia inoculation treatments on total parasite load in a model with gall count as the response variable and nitrogen treatment, rhizobia inoculation treatment, the nitrogen-by-inoculation interaction, and block as fixed effects (Table 1, Figure 1A). Because uninoculated, nodule-free plants could only gain nitrogen from the provided fertilizer, we compared uninoculated R-plants and inoculated R+ plants (combining the four rhizobia strains together) as a fixed effect to test the effect of mutualism establishment (inoculation) on parasite load. When using inoculation as a fixed effect, we fit rhizobia strain as a nested random effect, as the four strains formed a nested structure within the inoculated R+ group, creating non-independence within the broader inoculation effect. Our tests were robust to asymmetric sample sizes, but we acknowledge that they are relatively underpowered because there were many more inoculated (127) vs. uninoculated nodule-free (20) plants. We also subset the data for more symmetric comparison of 20 uninoculated plants to a randomly chosen focal strain consisting of 30 plants (USDA1021) with qualitatively similar results (Table S11; Table S12).

**Table 1:**
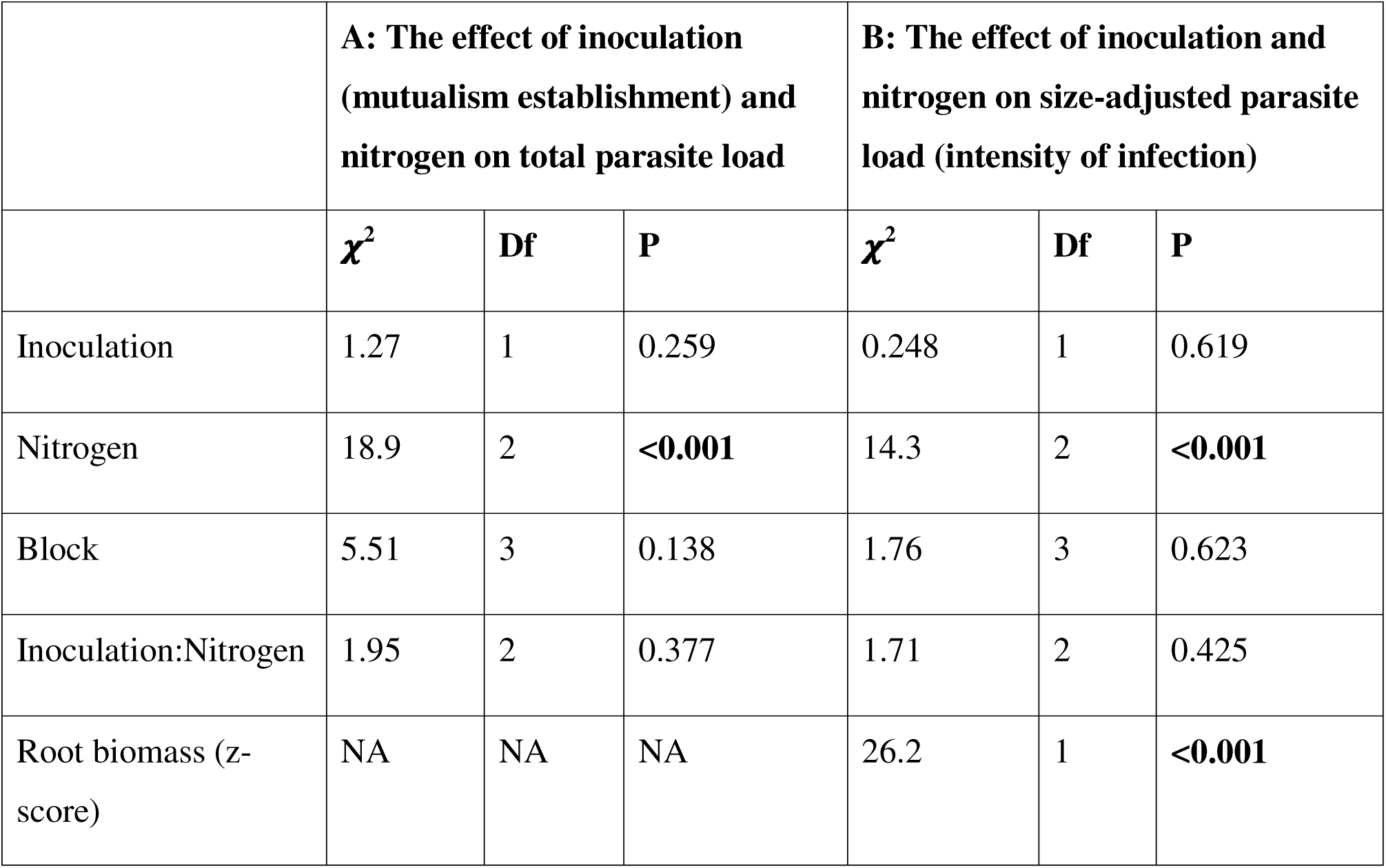
The effect of nitrogen and inoculation treatments on (A) parasite load and (B) size-adjusted parasite load, or intensity of infection. NA refers to not applicable, as only the second model includes root biomass to adjust for size.

**Figure 1:**
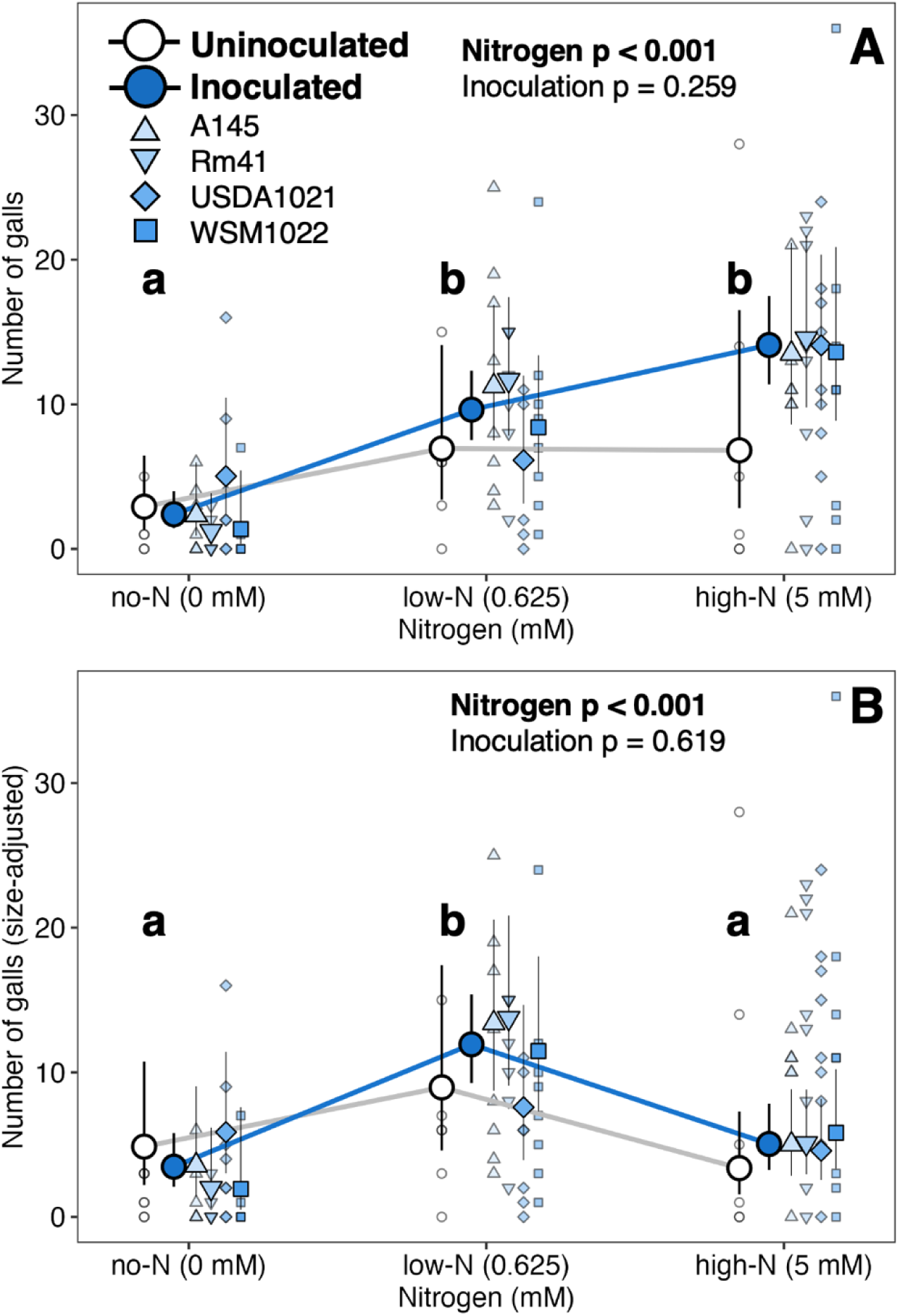
The effect of nitrogen and nodulation treatments on A) total parasite load and B) size-adjusted parasite load (infection intensity). Plants were inoculated with one of four different rhizobia strains which formed nodules; uninoculated plants were largely nodule-free. For panels, smaller transparent points are raw data, while the larger opaque points are the estimated marginal means. Error bars on the estimated marginal means show the 95% confidence interval. See Table 1 for ANOVA and Table S6 for all post-hoc tests.

We ran a second version of this first analysis that corrected for plant size by adding root biomass as an additional fixed effect (Table 1B, Figure 1B). We did so because greater nitrogen availability increased total biomass and nematodes formed more galls on larger plants (Figure S3, Table S3, Figure S9). This size-adjusted analysis (Figure 1B) more appropriately allows us to compare parasite load across our nitrogen treatments to assess the impact of nitrogen resources on parasite susceptibility. We view size-adjusted parasite load (hereafter referred to as the intensity of parasite infection) as a more appropriate estimate of the relative cost of infection. We did not directly measure host fitness costs of infection in this experiment.

Second, we tested for an association between non-nitrogen-fixing nodules and parasite load. Because these nodules did not provide nitrogen, a significant relationship between the two would support the mutualism establishment hypothesis. We standardized the number of non-N-fixing and N-fixing nodules within each nitrogen treatment using a z-score (a mean of zero and a standard deviation of one) to make comparisons across nitrogen treatments. We standardized nodule counts within each treatment because nodulation varies widely depending on nitrogen availability (Van Schreven 1959; Weese *et al*. 2015). We included nitrogen treatment, block, the number of non-N-fixing nodules, the number of N-fixing nodules, the interaction between nitrogen treatment and the number of non-N-fixing nodules, and the interaction between nitrogen treatment and the number of N-fixing nodules as fixed effects.

Finally, we tested whether the effect of nitrogen on intensity of parasite infection was explained by nitrogen assimilation into plant shoot tissues. We converted shoot nitrogen and carbon concentrations to z-scores within each nitrogen treatment by standardizing to a mean of zero and a standard deviation of one, after which they were incorporated as fixed effects for analysis. Nitrogen fertilizer treatment, nitrogen concentration, carbon concentration, root biomass, rhizobia strain, and block were included as fixed effects.

We confirmed all residual errors as normally distributed, homoscedastic, and properly dispersed using the DHARMa package (Hartig 2025). Post-hoc Tukey tests and least-squares means and confidence intervals were calculated with emmeans (Lenth 2023). All figures were created in ggplot2 (Wickham 2016).

## Results

### Mutualist-provided resources, not mutualism establishment, were correlated with increased parasite load

When we tested for an effect of our nitrogen and rhizobia inoculation treatments on total parasite load, we found a significant effect of nitrogen on total parasite load (p<0.001, Figure 1A, Table 1). Increasing nitrogen increased parasite load from no-N to low- and high-N (Figure 1A, Table S6). This effect was qualitatively similar across inoculated and uninoculated plants, and across rhizobia strains (Table 1, Table S5).

When we corrected for plant size to test for an effect of nitrogen and rhizobia inoculation treatments on the intensity of parasite infection, we found a significant effect of nitrogen on the intensity of parasite infection (size-adjusted parasite load) (p<0.001, Figure 1B, Table 1). Larger plants had more parasites overall, such that size-adjusted parasite load peaked at low-N, with no-N and high-N plants having similarly low intensities of infection (Figure S9, Figure 1A, Table S6).

In both of these tests, we did not find a significant effect of inoculation (mutualism establishment) on total or size-adjusted parasite load (Figure 1, Table 1). Inoculated and uninoculated plants did not significantly differ in total or size-adjusted parasite load (Figure 1, Table 1). Rhizobia strain had no significant effect on total or size-adjusted parasite load (Figure 1, Table S5). In fact, strains did not differ significantly in nodule number, nodule weight, resulting plant biomass, or total shoot nitrogen, although they did differ in in their proportion of N-fixing nodules (Figure S1, Table S1). Although this main test was underpowered, in general, we observed little significant variation associated with mutualism establishment in measured host traits, especially compared to the strong effects of the nitrogen fertilizer treatment.

Because our comparison of uninoculated (R-) and inoculated (R+) plants was underpowered, we then tested for an effect of non-nitrogen-fixing and nitrogen-fixing nodules on parasite load in a second analysis. We found no significant effect of non-nitrogen-fixing nodule count on total parasite load (Figure 2B, Table 2). This suggests that in the absence of nitrogen fixation, nodulation does not increase parasite infection. This is consistent with the analysis above, in which we observed no difference in infection between inoculation treatments. We did observe a significant effect of nitrogen-fixing nodule count on total parasite load: more nitrogen-fixing nodules corresponded to a higher total parasite load (p<0.001, Figure 2A, Table 2). Even accounting for the significant effect of N-fixing nodules, no-N plants still had a lower total parasite load than low- and high-N plants (Figure 2A, Table S7). Total host parasite load was sensitive to differing amounts of nitrogen provided to hosts by both exogenous fertilizer and by rhizobia in nitrogen-fixing nodules.

**Figure 2:**
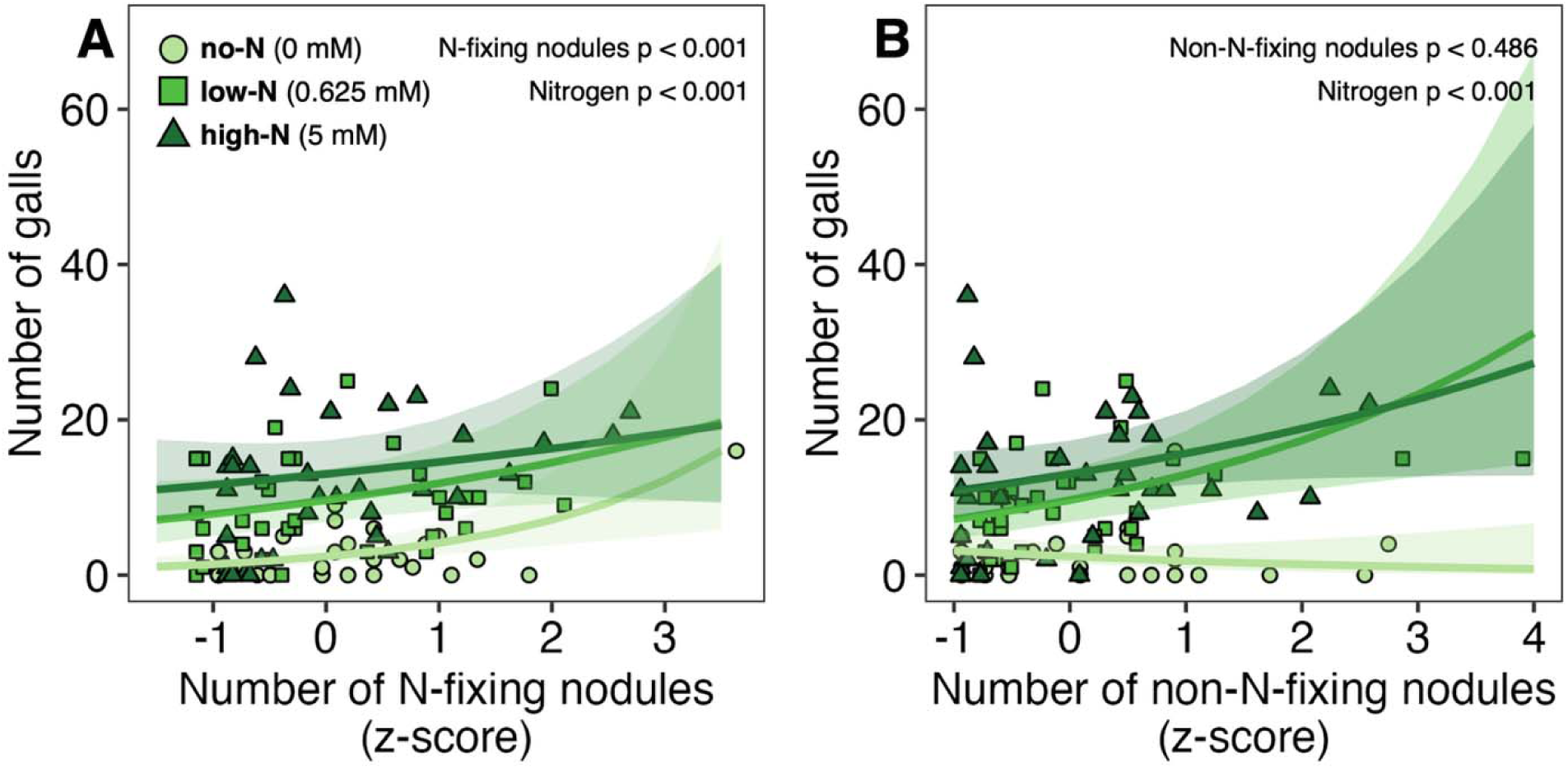
The relationship between nodulation (for N-fixing and non-N-fixing nodules) and gall number. N-fixing nodules had a significant effect on parasite load; non-N-fixing nodules did not. Different colors and shapes represent the different nitrogen fertilizer treatments, while each point consists of one plant sample. Nodule numbers (x-axis) were standardized to z-scores within each treatment.

**Table 2:**
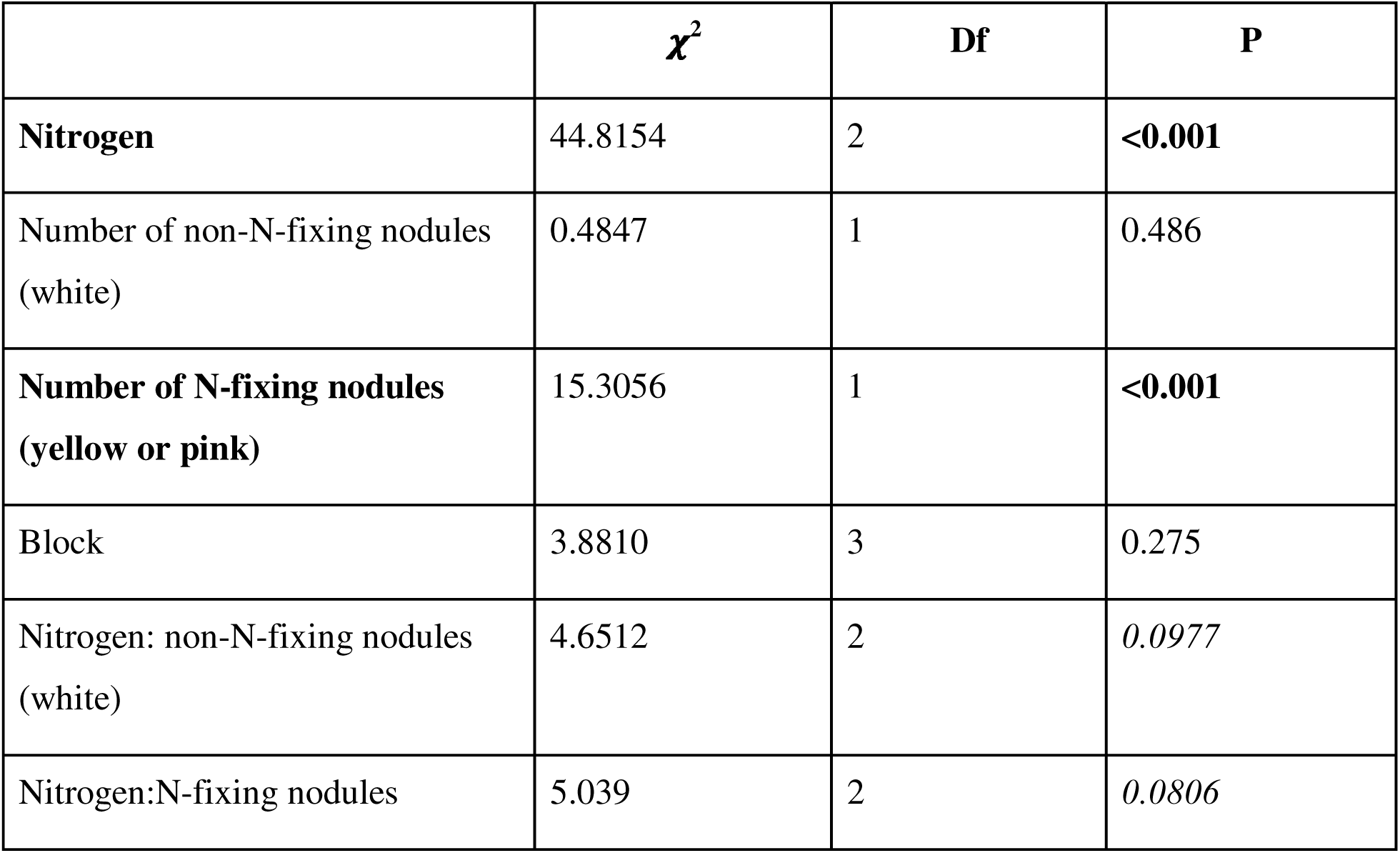
The relationship between nodulation (N-fixing and non-N-fixing) and gall number.

| | $\chi^2$ | Df | P |
| --- | --- | --- | --- |
| **Nitrogen** | 44.8154 | 2 | **<0.001** |
| Number of non-N-fixing nodules<br>(white) | 0.4847 | 1 | 0.486 |
| **Number of N-fixing nodules**<br>**(yellow or pink)** | 15.3056 | 1 | **<0.001** |
| Block | 3.8810 | 3 | 0.275 |
| Nitrogen: non-N-fixing nodules<br>(white) | 4.6512 | 2 | 0.0977 |
| Nitrogen:N-fixing nodules | 5.039 | 2 | 0.0806 |

### Mechanism of the nitrogen effect

Finally, when we tested whether the effect of nitrogen on intensity of parasite infection was explained by nitrogen assimilation into plant shoot tissues, we observed a significant effect of shoot tissue nitrogen concentration on intensity of parasite infection (p<0.001; Table 1). However, contrary to our initial expectation, higher nitrogen concentration led to a lower intensity of parasite infection (Table S8). Furthermore, significant differences in parasite infection between nitrogen treatments remained even after accounting for plant size and nitrogen concentration (p<0.001, Table 3). There was no relationship between carbon concentration and the intensity of parasite infection (p=0.181, Table 3).

**Table 3:**
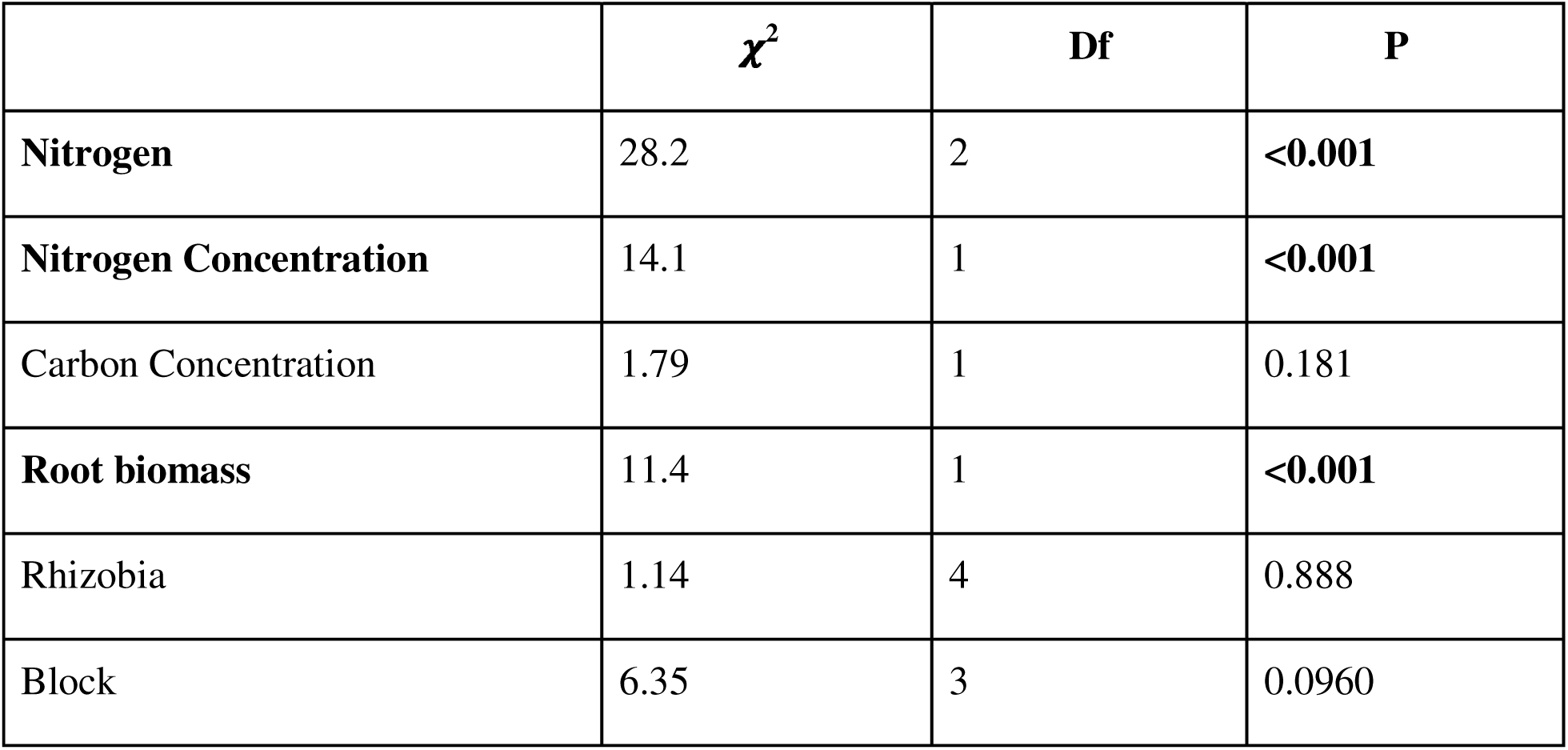
The relationship between nitrogen concentration and total parasite load.

## Discussion

In this study, we found that the resources provided by mutualistic symbionts were correlated with increased parasite infection. By contrast, mutualism establishment itself had little effect on parasitism here, despite the fact that legumes downregulate their local immune response when forming a symbiosis with rhizobia (Cao *et al*. 2017; Grundy *et al*. 2023).

However, the resource effect was more complicated than we anticipated, in multiple ways. First, the effect of nitrogen resources on the intensity of parasite infection was non-linear, peaking at an intermediate nitrogen level. Second, increased assimilated nitrogen in plants was correlated with *lower* infection. Third, the effect of nitrogen on parasite load was not fully explained by nitrogen assimilation into plant tissues, as a persistent effect of nitrogen treatment remained even after accounting for assimilation into tissues. Our results suggest that mutualist-provided resources may impact susceptibility to parasite infection through multiple potential avenues and physiological pathways, rather than by simply increasing attraction by parasites.

The results of this study suggest that nutritional mutualisms may increase susceptibility to infection, with infection risk potentially tied to the traded resource itself. If this is the case, then parasites and pathogens may be underappreciated constraints on the evolution of increased benefits in nutritional mutualisms. We conclude by exploring alternative hypotheses that might link mutualist-provided resources to the indirect costs of mutualism.

### Mutualist-provided resources increase parasite load, to a degree

Under the mutualist-provided resource hypothesis, we predicted that because mutualist-provided resources enrich host tissues, parasites would preferentially infect resource-rich hosts (Koricheva *et al*. 2009; Simonsen and Stinchcombe 2014).We expected this to manifest in our experiment as higher parasite loads at higher levels of nitrogen supplementation. As expected, the number of nitrogen-fixing nodules on a plant was positively correlated with increased parasite load (Figure 2A). We found, however, that nitrogen supplementation had a non-linear effect on parasite load. Although total parasite load increased from no-N to the low- and high-N fertilizer treatments, it leveled off between low- and high-N (Figure 1A). It is worth noting that while nitrogen provisioned through fertilizer and rhizobia fixation in nodules showed qualitatively similar patterns, we did not estimate the rate of nitrogen fixation in nodules and thus cannot directly compare the relationship of nitrogen provisioning to parasite load for these two nitrogen sources.

Furthermore, size-adjusted parasite load (i.e., the intensity of infection) peaked in the low-N treatment, and declined in the high-N treatment (Figure 1B). This pattern suggests that the relative cost of infection is highest at intermediate nitrogen levels, because although low- and high-N plants had similar numbers of parasites, low-N plants were more heavily infected given their overall size. Plants in the no-N treatment also had low parasite loads, but we suspect that this is simply because these plants were so small that they could not support any appreciable level of infection. In general, plant root size correlated with gall count, such that plants with larger roots had higher gall counts (Figure S9A). Our addition of a root biomass covariate to adjust for size allowed us to compare relative intensities of parasite infection (galls per unit of root mass).

### Nitrogen assimilation into tissues appears to be protective

However, shoot tissue nitrogen concentration was negatively correlated with parasite load (Figure 3, Table 3), the opposite of what would be expected if parasitic nematodes preferentially attacked nitrogen-rich hosts. This was surprising, given the extensive literature on preferential feeding by herbivores on nitrogen-rich tissue due to its better nutritional quality. Our experiment, however, was a no-choice experiment (parasites were not offered a choice between nitrogen-poor and nitrogen-rich hosts), which may explain why our results differ from these past studies.

**Figure 3:**
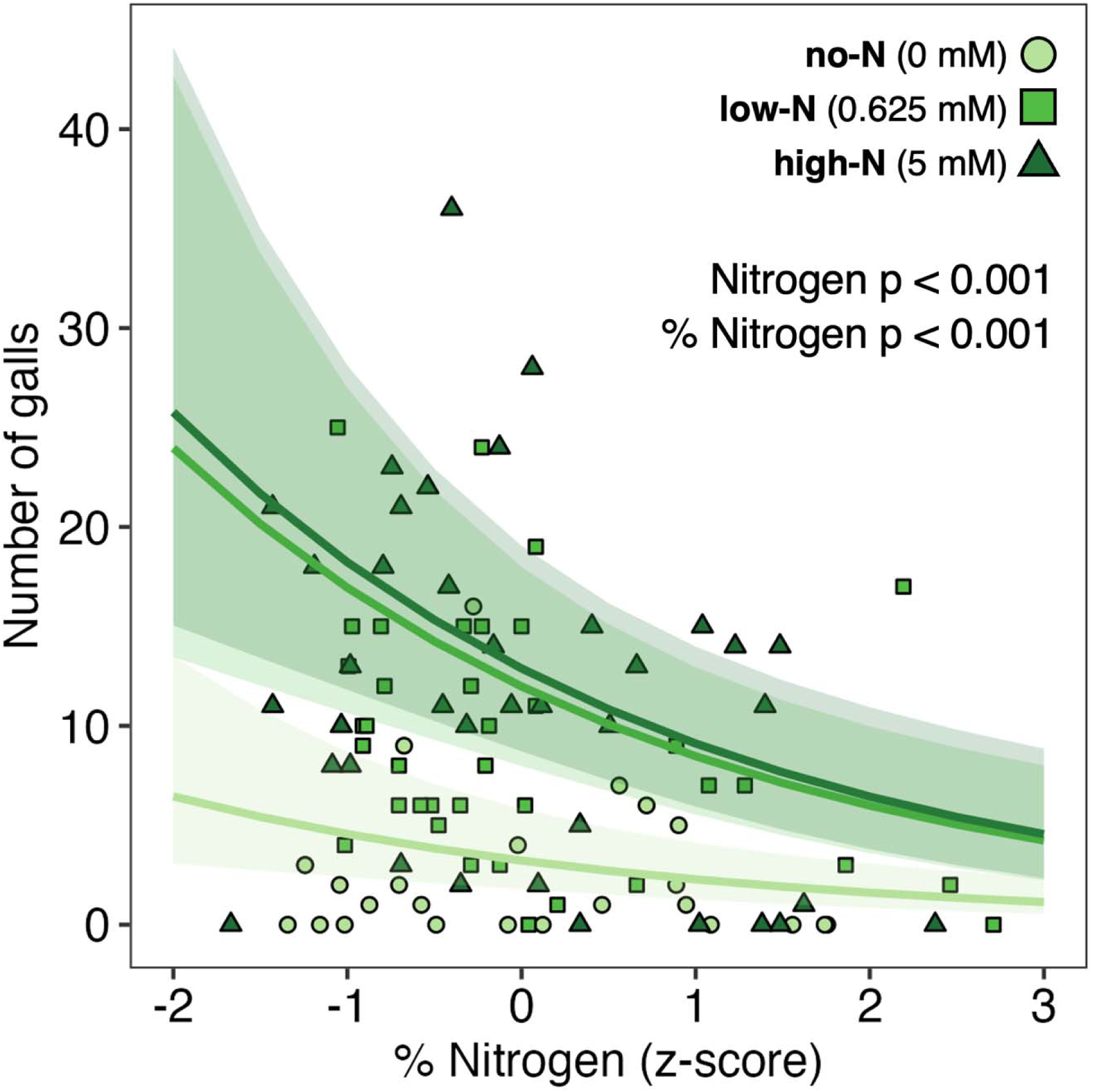
The relationship between shoot nitrogen concentration (x-axis) and total parasite load (y-axis). Different colors and shapes represent the different nitrogen fertilizer treatments, while each point consists of one plant sample. Both nitrogen fertilizer treatment as well as shoot nitrogen concentration affected the number of galls. Nitrogen concentration (x-axis) was z-scored within each treatment.

The negative relationship between nitrogen concentration and parasite load in our experiment suggests that nitrogen is protective and may play a role in plant defense responses. This is consistent with some other studies that have shown that nitrogen addition can decrease plant palatability in some contexts (Van Alstyne *et al*. 2009; Chen *et al*. 2010; Tomas *et al*. 2011). Hosts enriched with greater nitrogen resources may be able to mount stronger defenses, as higher nitrogen concentration in tissue may be indicative of nitrogen-rich host defense compounds. Many anti-herbivory defense compounds utilize nitrogen, such as alkaloids, cyanogenic glycosides, glucosinolates, and benzoxazinoids (Zenk and Juenger 2007). This might also account for the observed peak in the intensity of parasite infection at intermediate nitrogen levels (Figure 1B). This intermediate peak could represent a balance between parasite attraction to moderately N-rich plants and resistance to parasites by some extremely N-rich plants.

Alternatively, hosts enriched with greater nitrogen resources may also be more tolerant of parasites despite remaining susceptible. Larger size could reduce relative density and costs of parasites, while N-rich compounds or forms of dense storage may have deterred parasites outright. Plants allocate their resources differently depending on resource availability, such that defense may only become a priority over growth or reproduction at a certain threshold of resource richness (Herms and Mattson 1992; Züst and Agrawal 2017). Consistent with this hypothesis, plants in the high-N treatment allocated their additional nitrogen to either growing larger in size or increasing shoot nitrogen concentration (Figure S5).

### The resource effect is not fully explained by nitrogen assimilation into host tissues

Assimilated nitrogen does not fully explain the effect of nitrogen on parasite infection in our study. Even after accounting for the significant effect of shoot nitrogen concentration on total parasite load and intensity of infection, there remained a persistent difference in infection intensity between the intermediate- and high-nitrogen treatments (Figure 3). Nitrogen affects plants in many diverse, interlinked, and cascading ways which could impact parasite infection. It is an essential component of chlorophyll, which is necessary for photosynthesis. Various forms of nitrogen (nitrate and ammonium) function as signaling molecules within a complex network regulating uptake, metabolism, developmental phase change, and plant defense and immunity among other processes (Xuan *et al*. 2017; Vidal *et al*. 2020; Hancock 2020). Nitrate (NO ^-^) feeding—the form of our exogenous nitrogen fertilizer treatment—is thought to facilitate resistance through enhancing the production of defense signals, while ammonium (NH ^+^)—the product of bacteroid nitrogen fixation—may compromise defense through increased GABA nutrients for pathogens (Udvardi and Poole 2013; Mur *et al*. 2017). These differing forms of nitrogen may have affected our results, as rhizobia-provided ammonium might have contributed to higher parasite susceptibility than exogenous nitrate, which may have enhanced defense.

It is also possible that shoot nitrogen content, which we measured in this experiment, does not paint a complete picture of the relationship between plant nitrogen assimilation and root-knot nematode infection. Nematodes may have affected plant size and nitrogen content through their role as nutrient sinks: assimilated nitrogen may have been siphoned off to galls and surrounding root tissue, increasing root biomass through gall formation and potentially reducing shoot biomass and shoot nitrogen content (Figure S9). Therefore, though nitrogen content and assimilation appears to play a significant role in parasite load, considering the various fluxes of nitrogen and its uses to clarify the mechanisms of this correlation can contribute to a more complete understanding. Resources can also be differentially allocated to shoots, roots, leaves, or reproductive organs like seeds and fruit following optimal defense theory (Ohnmeiss and Baldwin 2000; Schultz *et al*. 2013). For instance, in our experiment, nitrogen may have driven increased investment in photosynthetic tissue (i.e., aboveground biomass) for plants growing in high-N (Figure S3, Figure S4).

### Little effect of mutualism establishment on parasite load

In this study, mutualism establishment alone did not have a significant effect on parasite load. We found no statistically significant differences in parasite load between host plants inoculated and uninoculated with rhizobia (Figure 1). One possible explanation for this was that our analysis was underpowered for the initial test between rhizobia-inoculated plants and uninoculated controls. However, we also showed in a second analysis that there was no relationship between non-nitrogen-fixing nodules and parasite load (Figure 2B), consistent with our conclusion that mutualism establishment (inoculation) alone did not increase parasite load. By contrast, there was an increase in parasite load with nitrogen-fixing nodules (Figure 2A), consistent with our conclusion that mutualist-provided resources may increase susceptibility to parasite infection in our system.

This finding is surprising because it is well known that legumes undergo significant changes in their immune system while forming symbioses with rhizobia. Nodulation is tightly regulated, with the host systematically downregulating its innate immune system to prevent plant defense responses upon detecting rhizobial Nod factors (Grundy *et al*. 2023). While we have high confidence about the effect of mutualist-provided nitrogen resources on parasite susceptibility, due to the underpowered nature of the first analysis, there remains the possibility of a weak to moderate effect of symbiosis or mutualism establishment on parasite susceptibility, Future research should interrogate why mutualism establishment alone seems to play a minor role in susceptibility to this particular parasite in the legume-rhizobia mutualism.

### Implications for mutualism ecology and evolution

In our experiment, nitrogen had a significant and complex effect on susceptibility to parasites. This finding has potentially wide-ranging implications because nitrogen is a common mutualist-provided resource. Rhizobia fix nitrogen for legumes; mycorrhizal fungi also form symbiotic associations with plant roots, benefiting plants’ ability to uptake nitrogen from the soil (Govindarajulu *et al*. 2005). Some cyanobacteria symbionts fix nitrogen for lichen (Zehr 2011). Similarly, algal turf communities fix nitrogen on coral reefs (Williams and Carpenter 1997). In ant-plant protection mutualisms and bird-plant pollinator mutualisms, animals will deposit nitrogen-rich droppings to benefit the plant (Defossez *et al*. 2011; Whelan *et al*. 2015; Pinkalski *et al*. 2018). Within nitrogen-limited environments, these mutualisms offset resource constraints, providing critical padding by increasing host and ecosystem nitrogen resource budgets (Berman-Frank *et al*. 2003; Guo *et al*. 2023). Nitrogen provisioning may be a common mechanism that increases susceptibility to antagonists across many different mutualisms.

In addition to mutualist-provided nitrogen, human activity in the age of the Anthropocene can also affect the ecology and outcomes of species interactions, as agricultural inputs contribute about 100 Tg of Nitrogen per year into soil through synthetic fertilizer, with additional organic inputs of manure and crop waste (Fowler *et al*. 2013). As such, many anthropogenic environments overturn the assumed norm of nitrogen limitation in natural environments. These human nitrogen inputs have dramatic downstream effects on ecosystems and species interactions, which are known to lead to potential breakdown of nitrogen-provisioning mutualisms (Johnson 1993; Weese *et al*. 2015). Moving forward, our data suggests that it would be worth investigating how anthropogenic nitrogen can cascade to other species interactions. Could increased susceptibility to parasites be an unexpected byproduct of resource input? The non-linear effect of nitrogen fertilization on parasite load in our results suggests that nitrogen may very quickly stop becoming the rate-limiting resource for host-parasite interactions. It has been proposed that the ecophysiology of carbon resource allocation can explain the ecology and evolution of mutualisms (Pringle 2016). Incorporating nitrogen or manipulating additional nutrients into a dynamic resource framework of production, exchange, and use can contribute to our understanding of indirect species interactions and ecosystem structure in a time of rapid anthropogenic change.

It is important to note that mutualist-provided resources increasing susceptibility to parasites does not necessarily mean that mutualists impose a net cost, resulting in antagonism (Bell *et al*. 2022). The interaction would still be considered mutualistic as long as the benefit outweighs the cost for both partners (Bronstein 2001b). The peak in size-adjusted parasite load at intermediate nitrogen levels may help to explain the wide range of effects of mutualisms on parasitism; while we focused on increasing susceptibility to parasites, many mutualisms do deter parasites (Clay 1988; González Teuber *et al*. 2014; Jandér 2015 p. 2023; Braquart-Varnier *et al*. 2015; Wang M *et al*. 2023). Nitrogen as a resource could both attract as well as defend against antagonists depending on the resource landscape, availability, and use. Future research should explore at what resource level the net benefits of mutualism are maximized, integrating the effect of resources on parasite infection as well as the direct benefit of resources.

Future research should explore the relative roles of mutualism establishment and mutualist-provided resources in driving susceptibility to parasites in other systems. Their relative importance likely varies across systems, because the host mounts a slightly different physiological response to each microorganism, and because different resources will likely differ in their cascading consequences for ecological interactions. Nevertheless, our study highlights the potential wide-ranging significance of nitrogen—with its massive biotic and anthropogenic inputs—for multi-species interactions and their outcomes. Finally, our results raise the intriguing possibility that nutritional mutualisms may inherently increase susceptibility to infection, because infection risk is tied directly to the traded resource. If this pattern extends beyond our study system to other nutritional mutualisms, then antagonists may be important overlooked agents of selection driving the evolution of increased benefits in nutritional mutualisms.

## Acknowledgements

We would like to thank Chigozie Ibe and Yoon Chang for data collection and their help in maintaining the experiment. We thank Ben Glass for his work on a previous version of the experiment. We also thank McCall Calvert, Addison Buxton-Martin, Chang-Yu Chang, Kim Gallagher, and Wood Lab members for their comments and feedback on the manuscript. The Plant Seminar group at the University of Pennsylvania provided helpful feedback on the experimental analysis and interpretation of results. We are grateful to Adrienne Gorny at North Carolina State University for *M. hapla* nematode lines, Nevin Young and the *Medicago* HapMap project for seeds, Qiulin Qin and Hongyan Zhu at the University of Kentucky for Rm41 and A145 rhizobia strains, the USDA for USDA1021 and WSM1022 rhizobia strains, and David Vann and the Instrumentation Core at the University of Pennsylvania’s Earth and Environmental Science Department for their assistance with elemental analysis. We are also grateful to Samara Gray and Kathryn Weber for their management of the greenhouses at the University of Pennsylvania. This work was funded by the National Science Foundation’s Division of Environmental Biology (grant no. 2118397) to Corlett Wood and a Penn Undergraduate Research Mentoring Program (PURM) fellowship award to Nova Meng.

## Literature Cited

Adler LS, Bronstein JL. 2004. Attracting Antagonists: Does Floral Nectar Increase Leaf Herbivory? Ecology 85: 1519–1526.

Barker DG, Pfaff T, Moreau D, et al. 2006. Growing M. truncatula: choice of substrates and growth conditions.

Bartlem DG, Jones MGK, Hammes UZ. 2014. Vascularization and nutrient delivery at root-knot nematode feeding sites in host roots. Journal of Experimental Botany 65: 1789–1798.

Batstone RT, Dutton EM, Wang D, Yang M, Frederickson ME. 2017. The evolution of symbiont preference traits in the model legume Medicago truncatula. New Phytologist 213: 1850–1861.

Bell CA, Magkourilou E, Urwin PE, Field KJ. 2022. Disruption of carbon for nutrient exchange between potato and arbuscular mycorrhizal fungi enhanced cyst nematode fitness and host pest tolerance. New Phytologist 234: 269–279.

Berman-Frank I, Lundgren P, Falkowski P. 2003. Nitrogen fixation and photosynthetic oxygen evolution in cyanobacteria. Research in Microbiology 154: 157–164.

Braquart-Varnier C, Altinli M, Pigeault R, et al. 2015. The Mutualistic Side of Wolbachia–Isopod Interactions: Wolbachia Mediated Protection Against Pathogenic Intracellular Bacteria. Frontiers in Microbiology 6.

Bronstein JL. 2001a. The exploitation of mutualisms. Ecology Letters 4: 277–287.

Bronstein JL. 2001b. The Costs of Mutualism. American Zoologist 41: 825–839.

Brooks M E, Kristensen K, Benthem K J, van, et al. 2017. glmmTMB Balances Speed and Flexibility Among Packages for Zero-inflated Generalized Linear Mixed Modeling. The R Journal 9: 378.

Burr AA, Woods KD, Cassidy ST, Wood CW. 2022. Priority effects alter the colonization success of a host-associated parasite and mutualist. Ecology 103: e3720.

Cao Y, Halane MK, Gassmann W, Stacey G. 2017. The Role of Plant Innate Immunity in the Legume-Rhizobium Symbiosis. Annual Review of Plant Biology 68: 535–561.

Castagnone-Sereno P, Danchin EGJ, Perfus-Barbeoch L, Abad P. 2013. Diversity and Evolution of Root-Knot Nematodes, Genus *Meloidogyne* : New Insights from the Genomic Era. Annual Review of Phytopathology 51: 203–220.

Catella SA, Olmsted CF, Markalanda SH, McFadden CJ, Wood CW, Kuebbing SE. 2022. A generalist nematode destabilises plant competition: no evidence for direct effects, but strong evidence for indirect effects on rhizobium abundance. New Phytologist 233: 2561–2572.

Charchar JM, Santo GS. 2009. Effect of soil temperature on the life cycle of Meloidogyne chitwoodi races 1 and 2 and M. hapla on Russet Burbank potato. Nematologia Brasileira 33:154–161.

Chaston J, Goodrich-Blair H. 2010. Common Trends in Mutualism Revealed by Model Associations Between Invertebrates and Bacteria. FEMS microbiology reviews 34: 41–58.

Chen Y, Olson DM, Ruberson JR. 2010. Effects of nitrogen fertilization on tritrophic interactions. Arthropod-Plant Interactions 4: 81–94.

Christian N, Whitaker BK, Clay K. 2015. Microbiomes: unifying animal and plant systems through the lens of community ecology theory. Frontiers in Microbiology 6.

Clay K. 1988. Fungal Endophytes of Grasses: A Defensive Mutualism between Plants and Fungi. Ecology 69: 10–16.

Cook DR. 1999. *Medicago truncatula* — a model in the making! Current Opinion in Plant Biology 2: 301–304.

Costa SR, Ng JLP, Mathesius U. 2021. Interaction of Symbiotic Rhizobia and Parasitic Root-Knot Nematodes in Legume Roots: From Molecular Regulation to Field Application. Molecular Plant-Microbe Interactions® 34: 470–490.

Crook MB, Lindsay DP, Biggs MB, et al. Griffitts JS, 2012. Rhizobial Plasmids That Cause Impaired Symbiotic Nitrogen Fixation and Enhanced Host Invasion. Mol Plant Microbe Interact 25(8): 1026–1033.

Damiani I, Baldacci-Cresp F, Hopkins J, et al. 2012. Plant genes involved in harbouring symbiotic rhizobia or pathogenic nematodes. New Phytologist 194: 511–522.

Defossez E, Djiéto-Lordon C, McKey D, Selosse M-A, Blatrix R. 2011. Plant-ants feed their host plant, but above all a fungal symbiont to recycle nitrogen. Proceedings of the Royal Society B: Biological Sciences 278: 1419–1426.

Evans, CS. 1991. J. B. Harborne (ED.) Methods in plant biochemistry: 1. plant phenolics. Academic Press, New York and London, 1990. £65. Phytochem. Anal., 2, 48–48.

Faeth SH, Fagan WF. 2002. Fungal Endophytes: Common Host Plant Symbionts but Uncommon Mutualists1. Integrative and Comparative Biology 42: 360–368.

Ferguson BJ, Mens C, Hastwell AH, et al. 2019. Legume nodulation: The host controls the party. Plant, Cell & Environment 42: 41–51.

Fischer HM. 1994. Genetic regulation of nitrogen fixation in rhizobia. Microbiological Reviews 58: 352–386.

Forrester NJ, Rebolleda-Gómez M, Sachs JL, Ashman TL, 2020. Polyploid Plants Obtain Greater Fitness Benefits from a Nutrient Acquisition Mutualism. New Phytologist 227, no. 3 : 944–54.

Fowler D, Coyle M, Skiba U, et al. 2013. The global nitrogen cycle in the twenty-first century. Philosophical Transactions of the Royal Society B: Biological Sciences 368: 20130164.

Frederickson ME, Gordon DM. 2007. The devil to pay: a cost of mutualism with *Myrmelachista schumanni* ants in ‘devil’s gardens’ is increased herbivory on *Duroia hirsuta* trees. Proceedings of the Royal Society B: Biological Sciences 274: 1117–1123.

Fuess LE, Palacio-Castro AM, Butler CC, Baker AC, Mydlarz LD. 2020. Increased Algal Symbiont Density Reduces Host Immunity in a Threatened Caribbean Coral Species, Orbicella faveolata. Frontiers in Ecology and Evolution 8.

Gelman A, Hill J. 2007. Data Analysis Using Regression and Multilevel/Hierarchical Models. Cambridge University Press.

González Teuber M, Kaltenpoth M, Boland W. 2014. Mutualistic ants as an indirect defence against leaf pathogens. New Phytologist 202: 640–650.

Gourion B, Berrabah F, Ratet P, Stacey G. 2015. Rhizobium–legume symbioses: the crucial role of plant immunity. Trends in Plant Science 20: 186–194.

Govindarajulu M, Pfeffer PE, Jin H, et al. 2005. Nitrogen transfer in the arbuscular mycorrhizal symbiosis. Nature 435: 819–823.

Grundy EB, Gresshoff PM, Su H, Ferguson BJ. 2023. Legumes Regulate Symbiosis with Rhizobia via Their Innate Immune System. International Journal of Molecular Sciences 24: 2800.

Guo Y, Fudali S, Gimeno J, et al. 2017. Networks Underpinning Symbiosis Revealed Through Cross-Species eQTL Mapping. Genetics 206: 2175–2184.

Guo K, Yang J, Yu N, Luo L, Wang E. 2023. Biological nitrogen fixation in cereal crops: Progress, strategies, and perspectives. Plant Communications 4: 100499.

Hancock JT. 2020. Nitric Oxide Signaling in Plants. Plants 9: 1550.

Harrison TL, Wood CW, Heath KD, Stinchcombe JR. 2017. Geographically structured genetic variation in the Medicago lupulina–Ensifer mutualism. Evolution 71: 1787–1801.

Hartig, F. 2025. DHARMa: Residual Diagnostics for Hierarchical (Multi-Level / Mixed) Regression Models. R package version 0.4.7. http://florianhartig.github.io/DHARMa/

Hartley SE, Gange AC. 2009. Impacts of Plant Symbiotic Fungi on Insect Herbivores: Mutualism in a Multitrophic Context. Annual Review of Entomology 54: 323–342.

Herms DA, Mattson WJ. 1992. The Dilemma of Plants: To Grow or Defend. The Quarterly Review of Biology 67: 283–335.

Herrera CM, Medrano M, Rey PJ, et al. 2002. Interaction of pollinators and herbivores on plant fitness suggests a pathway for correlated evolution of mutualism- and antagonism-related traits. Proceedings of the National Academy of Sciences 99: 16823–16828.

Irmer S, Podzun N, Langel D, et al. 2015. New aspect of plant–rhizobia interaction: Alkaloid biosynthesis in Crotalaria depends on nodulation. Proceedings of the National Academy of Sciences of the United States of America 112: 4164–4169.

Irwin RE, Bronstein JL, Manson JS, Richardson L. 2010. Nectar Robbing: Ecological and Evolutionary Perspectives. Annual Review of Ecology, Evolution, and Systematics 41: 271–292.

Jandér KC. 2015. Indirect mutualism: ants protect fig seeds and pollen dispersers from parasites. Ecological Entomology 40: 500–510.

Janzen, DH. 1975. *Pseudomyrmex nigropilosa* : A Parasite of a Mutualism. Science, 188, 936–937.

Janzen DH, Boucher D. 1985. The natural history of mutualisms In: The Biology of Mutualism. 40–99.

Johnson NC. 1993. Can Fertilization of Soil Select Less Mutualistic Mycorrhizae? Ecological Applications 3: 749–757.

Johnson CA, Bronstein JL. 2019. Coexistence and competitive exclusion in mutualism. Ecology 100: e02708.

Johnstone RA, Bshary R. 2002. From parasitism to mutualism: partner control in asymmetric interactions. Ecology Letters 5: 634–639.

Jung SC, Martinez-Medina A, Lopez-Raez JA, Pozo MJ. 2012. Mycorrhiza-induced resistance and priming of plant defenses. Journal of Chemical Ecology 38: 651–664.

Kandouh BH, Hasan AE, Al-Hakeem AMA, Aridh ZHJ. A Complete Safe and Cost-Effective Method For Staining Root-Knot Nematodes *Meloidogyne* spp.

Kearsley JVS, Sather LM, Finan TM. 2024. Sinorhizobium (Ensifer) meliloti. Trends in Microbiology 32: 516–518.

Klepzig KD, Adams AS, Handelsman J, Raffa KF. 2009. Symbioses: A Key Driver of Insect Physiological Processes, Ecological Interactions, Evolutionary Diversification, and Impacts on Humans*. Environmental Entomology 38: 67–77.

Koricheva J, Gange AC, Jones T. 2009. Effects of mycorrhizal fungi on insect herbivores: a meta analysis. Ecology 90: 2088–2097.

Lenth, RV. 2023. emmeans: Estimated Marginal Means, aka Least-Squares Means. R package version 1.8.7.

Lindström K, Mousavi SA. 2020. Effectiveness of nitrogen fixation in rhizobia. Microbial Biotechnology 13: 1314–1335.

Little AE, Currie CR. 2009. Parasites may help stabilize cooperative relationships. BMC Evolutionary Biology 9: 124.

Mansfield KM, Gilmore TD. 2019. Innate immunity and cnidarian-Symbiodiniaceae mutualism. Developmental & Comparative Immunology 90: 199–209.

Masson-Boivin C, Sachs JL. 2018. Symbiotic nitrogen fixation by rhizobia-the roots of a success story. Current Opinion in Plant Biology 44: 7–15.

Meehan CJ, Olson EJ, Reudink MW, Kyser TK, Curry RL. 2009. Herbivory in a spider through exploitation of an ant–plant mutualism. Current Biology 19: R892–R893.

Miller R m.1993. Nontarget and ecological effects of transgenically altered disease resistance in crops -possible effects on the mycorrhizal symbiosis. Molecular Ecology 2: 327–335.

Moland E, Jones GP. 2004. Experimental confirmation of aggressive mimicry by a coral reef fish. Oecologia 140: 676–683.

Moreau D, Voisin A-S, Salon C, Munier-Jolain N. 2008. The model symbiotic association between Medicago truncatula cv. Jemalong and Rhizobium meliloti strain 2011 leads to N-stressed plants when symbiotic N2 fixation is the main N source for plant growth. Journal of Experimental Botany 59: 3509–3522.

Mur LAJ, Simpson C, Kumari A, Gupta AK, Gupta KJ. 2017. Moving nitrogen to the centre of plant defence against pathogens. Annals of Botany 119: 703–709.

Ness JH. 2006. A mutualism’s indirect costs: the most aggressive plant bodyguards also deter pollinators. Oikos 113: 506–514.

Ohnmeiss TE, Baldwin IT. 2000. Optimal Defense Theory Predicts the Ontogeny of an Induced Nicotine Defense. Ecology 81: 1765–1783.

Opitz MW, Díaz-Manzano FE, Ruiz-Ferrer V, et al. Wieczorek K, 2024. “The other side of the coin: systemic effects of Serendipita indica root colonization on development of sedentary plant-parasitic nematodes in Arabidopsis thaliana.” Planta. 2024;259(5):121.

Ott T, Van Dongen JT, Günther C, et al. 2005. Symbiotic Leghemoglobins Are Crucial for Nitrogen Fixation in Legume Root Nodules but Not for General Plant Growth and Development. Current Biology 15: 531–535.

Patil, J, Miller AJ, Gaur HS, 2013. “Effect of Nitrogen Supply Form on the Invasion of Rice Roots by the Root-Knot Nematode, Meloidogyne Graminicola.” Nematology 15, no. 4: 483–92.

Pinkalski C, Jensen KV, Damgaard C, Offenberg J. 2018. Foliar uptake of nitrogen from ant faecal droplets: An overlooked service to ant plants (R McCulley, Ed.). Journal of Ecology 106: 289–295.

Poll J, Marhan S, Haase S, Hallmann J, Kandeler E, Ruess L, 2007.. “Low Amounts of Herbivory by Root-Knot Nematodes Affect Microbial Community Dynamics and Carbon Allocation in the Rhizosphere.” FEMS Microbiology Ecology 62, no. 3:268–79.

Prévitali T, Rouault M, Pichereaux C, Gourion B. 2025. Lotus resistance against Ralstonia is enhanced by Mesorhizobium and does not impair mutualism. New Phytologist 245: 1249–1262.

Pringle EG. 2016. Integrating plant carbon dynamics with mutualism ecology. New Phytologist 210: 71–75.

Raine NE, Gammans N, Macfadyen IJ, Scrivner GK, Stone GN. 2004. Guards and thieves: antagonistic interactions between two ant species coexisting on the same ant plant. Ecological Entomology 29: 345–352.

Regus J., Quides K, O’Neill MR, et al. Sachs JL, 2017. “Cell Autonomous Sanctions in Legumes Target Ineffective Rhizobia in Nodules with Mixed Infections.” American Journal of Botany 104, no. 9: 1299–312.

Rice MM, Baldwin DG, Fischer JN, Fuchs C, Burkepile DE. 2021. Complex interactions with nutrients and sediment alter the effects of predation on a reef building coral. Marine Ecology 42: e12670.

Richardson DM, Allsopp N, D’Antonio CM, Milton SJ, Rejmánek M. 2000. Plant invasions – the role of mutualisms. Biological Reviews of the Cambridge Philosophical Society 75: 65–93.

Rose RJ. 2008. Medicago truncatula as a model for understanding plant interactions with other organisms, plant development and stress biology: past, present and future. Functional plant biology: FPB 35: 253–264.

Saikkonen K, Saari S, Helander M. 2010. Defensive mutualism between plants and endophytic fungi? Fungal Diversity 41: 101–113.

Salem H, Onchuru TO, Bauer E, Kaltenpoth M. 2015. Symbiont transmission entails the risk of parasite infection. Biology Letters 11: 20150840.

Schmitt RJ, Holbrook SJ. 2003. Mutualism can mediate competition and promote coexistence. Ecology Letters 6: 898–902.

Schultz JC, Appel HM, Ferrieri A, Arnold TM. 2013. Flexible resource allocation during plant defense responses. Frontiers in Plant Science 4.

Shanmugam KT, O’Gara F, Andersen K, Valentine RC. 1978. Biological Nitrogen Fixation. Annual Review of Plant Biology 29: 263–276.

Simonsen AK, Stinchcombe JR. 2014. Herbivory eliminates fitness costs of mutualism exploiters. New Phytologist 202: 651–661.

Smith, DC. 1991. Foreword. In L. Margulis and R. Fester (eds.), Symbiosis as a Source of Evolutionary Innovation, pp. ix–x. Cambridge, MA: The MIT Press.

Tajima R, Lee ON, Abe J, Lu A, Morita S, 2007. “Nitrogen-Fixing Activity of Root Nodules in Relation to Their Size in Peanut (Arachis Hypogaea L.).” Plant Production Science 10, no. 4: 423–29. 10.1626/pps.10.423.

Tomas F, Abbott JM, Steinberg C, Balk M, Williams SL, Stachowicz JJ. 2011. Plant genotype and nitrogen loading influence seagrass productivity, biochemistry, and plant–herbivore interactions. Ecology 92: 1807–1817.

Toth R, Toth D, Starke D, Smith DR. 1990. Vesicular–arbuscular mycorrhizal colonization in Zea mays affected by breeding for resistance to fungal pathogens. Canadian Journal of Botany 68: 1039–1044.

Traveset A, Richardson DM. 2014. Mutualistic Interactions and Biological Invasions. Annual Review of Ecology, Evolution, and Systematics 45: 89–113.

Truong NM, Nguyen C-N, Abad P, Quentin M, Favery B. 2015. Chapter Twelve - Function of Root-Knot Nematode Effectors and Their Targets in Plant Parasitism In: Escobar C, Fenoll C, eds. Plant Nematode Interactions. Advances in Botanical Research. Academic Press, 293–324.

Udvardi M, Poole PS. 2013. Transport and Metabolism in Legume-Rhizobia Symbioses. Annual Review of Plant Biology 64: 781–805.

Unger S, Habermann FM, Schenke K, Jongen M. 2021. Arbuscular Mycorrhizal Fungi and Nutrition Determine the Outcome of Competition Between Lolium multiflorum and Trifolium subterraneum. Frontiers in Plant Science 12.

Van Alstyne KL, Pelletreau KN, Kirby A. 2009. Nutritional preferences override chemical defenses in determining food choice by a generalist herbivore, Littorina sitkana. Journal of Experimental Marine Biology and Ecology 379: 85–91.

Van Schreven DA. 1959. Effects of added sugars and nitrogen on nodulation of legumes. Plant and Soil 11: 93–112.

Vidal EA, Alvarez JM, Araus V, et al. 2020. Nitrate in 2020: Thirty Years from Transport to Signaling Networks. The Plant Cell 32: 2094–2119.

Wang LX, Wang LL, Tan Q, et al. 2016. Efficient Inactivation of Symbiotic Nitrogen Fixation Related Genes in Lotus japonicus Using CRISPR-Cas9. Frontiers in Plant Science 7.

Wang LL, Rubio MC, Xin X, et al. 2019. CRISPR/Cas9 knockout of leghemoglobin genes in *Lotus japonicus* uncovers their synergistic roles in symbiotic nitrogen fixation. New Phytologist 224: 818–832.

Wang, L, Chen X, Yan X, Wang C, Guan P, Tang Z, 2023. “A Response of Biomass and Nutrient Allocation to the Combined Effects of Soil Nutrient, Arbuscular Mycorrhizal, and Root-Knot Nematode in Cherry Tomato.” Frontiers in Ecology and Evolution 11:1106122.

Wang M, Wang Z, Guo M, Qu L, Biere A. 2023. Effects of arbuscular mycorrhizal fungi on plant growth and herbivore infestation depend on availability of soil water and nutrients. Frontiers in Plant Science 14: 1101932.

Wang Q, Liu J, Li H, et al. 2018. Nodule-Specific Cysteine-Rich Peptides Negatively Regulate Nitrogen-Fixing Symbiosis in a Strain-Specific Manner in *Medicago truncatula*. Molecular Plant-Microbe Interactions® 31: 240–248.

Weese DJ, Heath KD, Dentinger BTM, Lau JA. 2015. Long-term nitrogen addition causes the evolution of less-cooperative mutualists: NITROGEN ADDITION DESTABILIZES MUTUALISM. Evolution 69: 631–642.

Wendlandt CE, Gano-Cohen KA, Stokes PJN, et al. Sachs JL, 2022. “Wild Legumes Maintain Beneficial Soil Rhizobia Populations despite Decades of Nitrogen Deposition.” Oecologia 198, no. 2: 419–30.

Whelan CJ, Şekercioğlu ÇH, Wenny DG. 2015. Why birds matter: from economic ornithology to ecosystem services. Journal of Ornithology 156: 227–238.

Wickham H. 2016. ggplot2. Cham: Springer International Publishing.

Williams SL, Carpenter RC. 1997. Grazing effects on nitrogen fixation in coral reef algal turfs. Marine Biology 130: 223–231.

Willig JJ, Sonneveld D, van Steenbrugge JJM, et al. Sterken, MG, 2023. “From Root to Shoot: Quantifying Nematode Tolerance in Arabidopsis Thaliana by High-Throughput Phenotyping of Plant Development.” Journal of Experimental Botany 74, no. 18 5487–99.

Wood CW, Pilkington BL, Vaidya P, Biel C, Stinchcombe JR. 2018. Genetic conflict with a parasitic nematode disrupts the legume–rhizobia mutualism. Evolution Letters 2: 233–245.

Xuan W, Beeckman T, Xu G. 2017. Plant nitrogen nutrition: sensing and signaling. Current Opinion in Plant Biology 39: 57–65.

Yin W, Zhou H., Wu M., et al. Ding J. 2025. “Reassociation of Specialist Herbivores with an Invasive Plant Selects for Reduced Allocation to Soil Mutualists.” New Phytologist. 10.1111/nph.70647.

Yu L, Zhang W, Geng Y, Liu K, Shao X. 2022. Cooperation With Arbuscular Mycorrhizal Fungi Increases Plant Nutrient Uptake and Improves Defenses Against Insects. Frontiers in Ecology and Evolution 10.

Zehr JP. 2011. Nitrogen fixation by marine cyanobacteria. Trends in Microbiology 19: 162–173.

Zenk MH, Juenger M. 2007. Evolution and current status of the phytochemistry of nitrogenous compounds. Phytochemistry 68: 2757–2772.

Züst T, Agrawal AA. 2017. Trade-Offs Between Plant Growth and Defense Against Insect Herbivory: An Emerging Mechanistic Synthesis. Annual Review of Plant Biology 68: 513–534.

