## Supplement for "Mutualist-Provided Resources Increase Susceptibility to Parasites"

### **Supplementary Materials**

### **Figures & Tables**

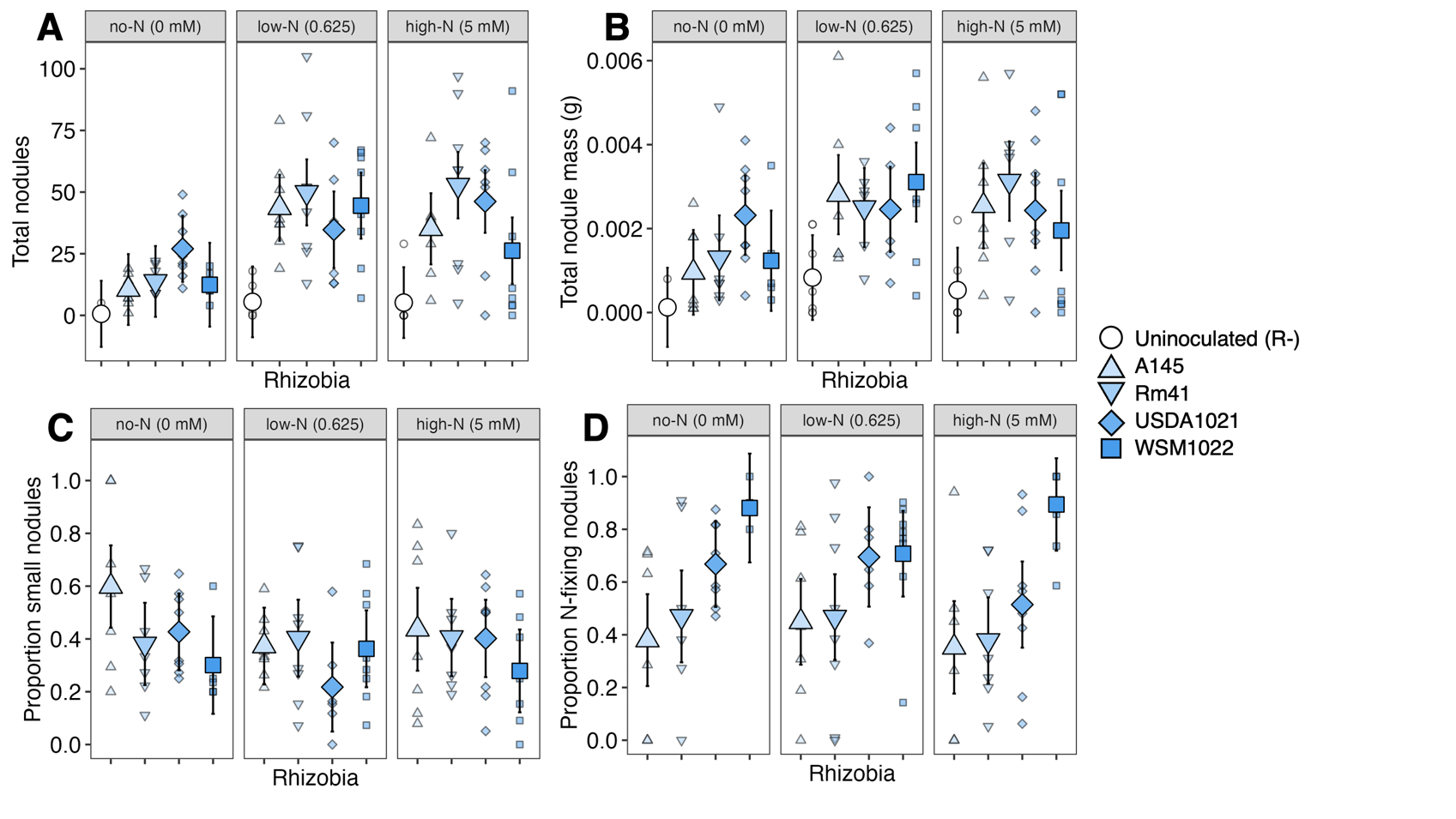

**Figure S1:** Nodule characteristics by rhizobia strain. For **A** and **B**, R- contains all plants, irrespective of contamination status. For all panels, smaller transparent points are raw data, while the larger opaque points are the estimated marginal means. Error bars on the estimated marginal means show the 95% confidence interval.

**A.** The effect of rhizobia strain and nitrogen treatment on the total number of nodules. Rhizobia strain and nitrogen both had an effect on the total number of nodules (p<0.001 for both); the interaction effect for rhizobia strain and nitrogen was marginally significant (p=0.0768). All contaminated uninoculated plants were included.

**B.** The effect of rhizobia strain and nitrogen treatment on total nodule mass. Rhizobia strain and nitrogen both had an effect on total nodule mass (p<0.001 for both); block had a marginally significant effect (p=0.0552). All contaminated uninoculated plants were included.

**C.** The effect of rhizobia strain and nitrogen treatment on proportion of small nodules. Rhizobia strain had a marginally significant effect (p=0.0847).

**D.** The effect of rhizobia strain and nitrogen treatment on proportion of N-fixing nodules. Rhizobia strain and block both had significant effects (p<0.001 and p=0.00231).

**Table S1:** The effect of rhizobia strains and nitrogen on nodule traits.

|  | **X^2^** | **Df** | **P** |
| --- | --- | --- | --- |
| A. **Effect of rhizobia strain and nitrogen treatment on total number of nodules** | | | |
| **Nitrogen** | 30.0 | 2 | **<0.001** |
| **Rhizobia** | 46.5 | 4 | **<0.001** |
| Block | 1.39 | 3 | 0.708 |
| Nitrogen:Rhizobia | 14.2 | 8 | 0.0768 |
| B. **Effect of rhizobia strain and nitrogen treatment on total nodule mass** | | | |
| **Nitrogen** | 14.9 | 2 | **<0.001** |
| **Rhizobia** | 31.4 | 4 | **<0.001** |
| Block | 7.60 | 3 | 0.0552 |
| Nitrogen:Rhizobia | 9.00 | 8 | 0.342 |
| C. **Effect of rhizobia strain and nitrogen treatment on proportion of small nodules** | | | |
| Nitrogen | 2.49 | 2 | 0.287 |
| Rhizobia | 6.63 | 3 | 0.0847 |
| Block | 0.311 | 3 | 0.958 |
| Nitrogen:Rhizobia | 6.86 | 6 | 0.334 |
| D. **Effect of rhizobia strain and nitrogen treatment on proportion of N-fixing nodules** | | | |
| Nitrogen | 1.16 | 2 | 0.559 |
| **Rhizobia** | 45.2 | 3 | **<0.001** |
| **Block** | 14.5 | 3 | **0.00231** |
| Nitrogen:Rhizobia | 5.87 | 6 | 0.438 |
| E. **Comparison of proportion of N-fixing nodules by rhizobia strain** | | | |
|  | **Ratio** | **SE** | **P** |
| A145 and Rm41 | -0.647 | 0.0690 | 0.916 |
| **A145 and USDA1021** | -3.31 | 0.0701 | **0.0078** |
| **A145 and WSM1022** | -6.01 | 0.0722 | **<0.001** |
| **Rm41 and USDA1021** | -2.70 | 0.0695 | **0.0421** |
| **Rm41 and WSM1022** | -5.44 | 0.0715 | **<0.001** |
| **USDA1021 and WSM1022** | -2.78 | 0.0725 | **0.0343** |

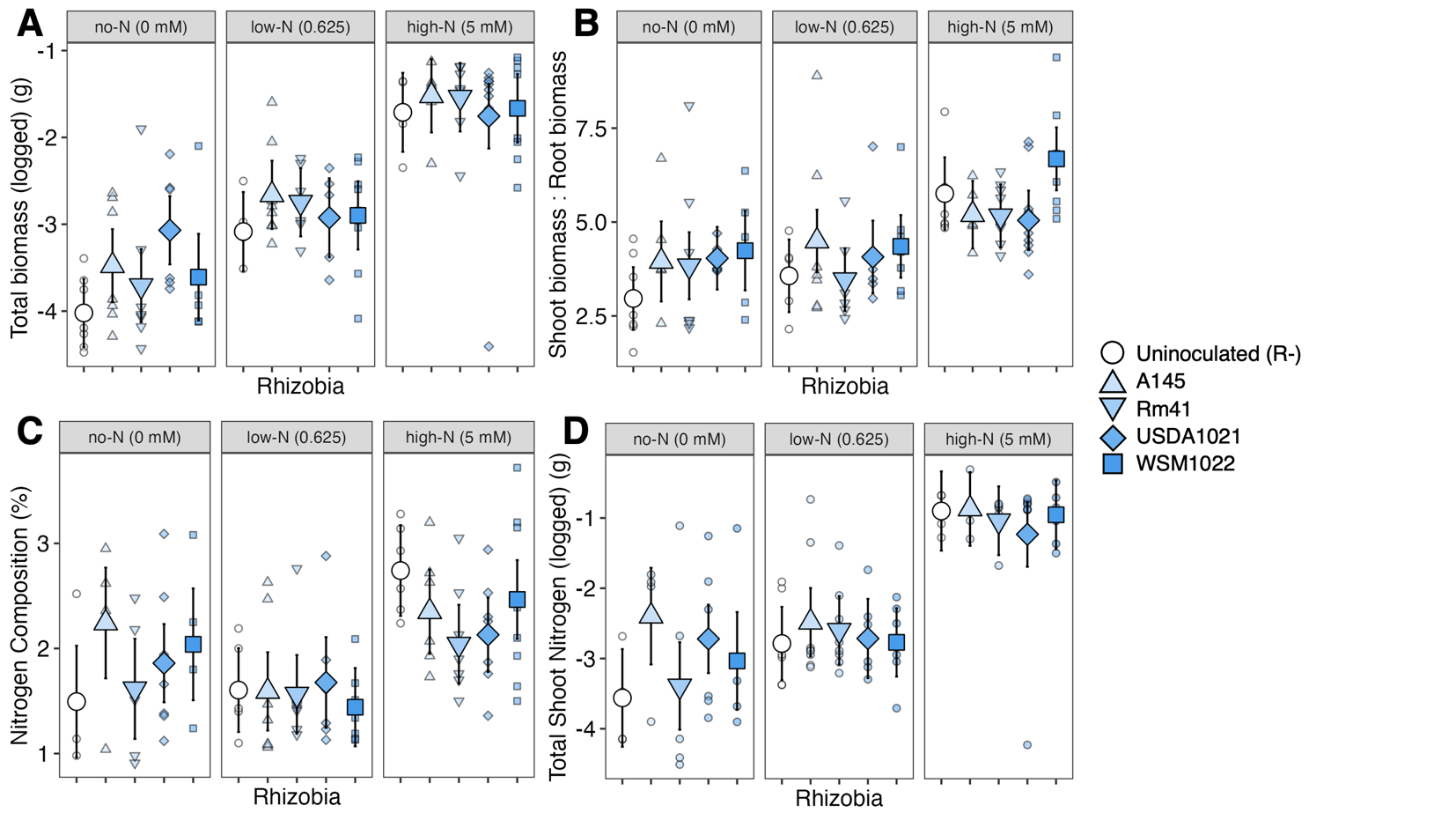

**Figure S2:** The effect of rhizobia strains on host plant resource budgets. For all panels, smaller transparent points are raw data, while the larger opaque points are the estimated marginal means. Error bars on the estimated marginal means show the 95% confidence interval.

**A.** Total biomass for rhizobia strains and nitrogen treatments. Nitrogen treatment had a significant effect on total biomass, but rhizobia strain did not. Contaminated uninoculated plants that formed 18 or more nodules were excluded.

**B.** Shoot:Root ratios for rhizobia strains and nitrogen treatments. Rhizobia strain and nitrogen both had significant effects on shoot:root ratios. Contaminated uninoculated plants that formed 18 or more nodules were excluded.

**C.** Shoot nitrogen concentration (%N) for rhizobia strains and nitrogen treatments. Nitrogen treatment had a significant effect on tissue concentration; rhizobia strain did not. Contaminated uninoculated plants that formed 18 or more nodules were excluded.

**D.** Logged total shoot nitrogen content for rhizobia strains and nitrogen treatments. Nitrogen treatment had a significant effect on total shoot nitrogen content; rhizobia strain did not. Contaminated uninoculated plants that formed 18 or more nodules were excluded.

**Table S2:** The effect of rhizobia strains on host plant resource budgets.

|  | **X ^2^** | **Df** | **P** |
| --- | --- | --- | --- |
| A. **Effect of rhizobia strain and nitrogen on total biomass (logged)** | | | |
| **Nitrogen** | 219 | 2 | **<0.001** |
| Rhizobia | 6.07 | 4 | 0.194 |
| Block | 2.79 | 3 | 0.426 |
| Nitrogen:Rhizobia | 8.64 | 8 | 0.374 |
| B. **Effect of rhizobia strain and nitrogen on shoot:root ratios** | | | |
| **Nitrogen** | 47.1 | 2 | **<0.001** |
| Rhizobia | 9.14 | 4 | 0.0578 |
| **Block** | 10.34 | 3 | **0.0159** |
| Nitrogen:Rhizobia | 9.18 | 8 | 0.328 |
| C. **Effect of rhizobia strain and nitrogen on shoot nitrogen concentration** | | | |
| **Nitrogen** | 39.1 | 2 | **<0.001** |
| Rhizobia | 3.86 | 4 | 0.425 |
| Block | 3.71 | 3 | 0.295 |
| Nitrogen:Rhizobia | 10.6 | 8 | 0.222 |
| D. **Effect of rhizobia strain and nitrogen on total shoot nitrogen** | | | |
| **Nitrogen** | 156 | 2 | **<0.001** |
| Rhizobia | 5.26 | 4 | 0.261 |
| Block | 4.61 | 3 | 0.203 |
| Nitrogen:Rhizobia | 7.42 | 8 | 0.492 |

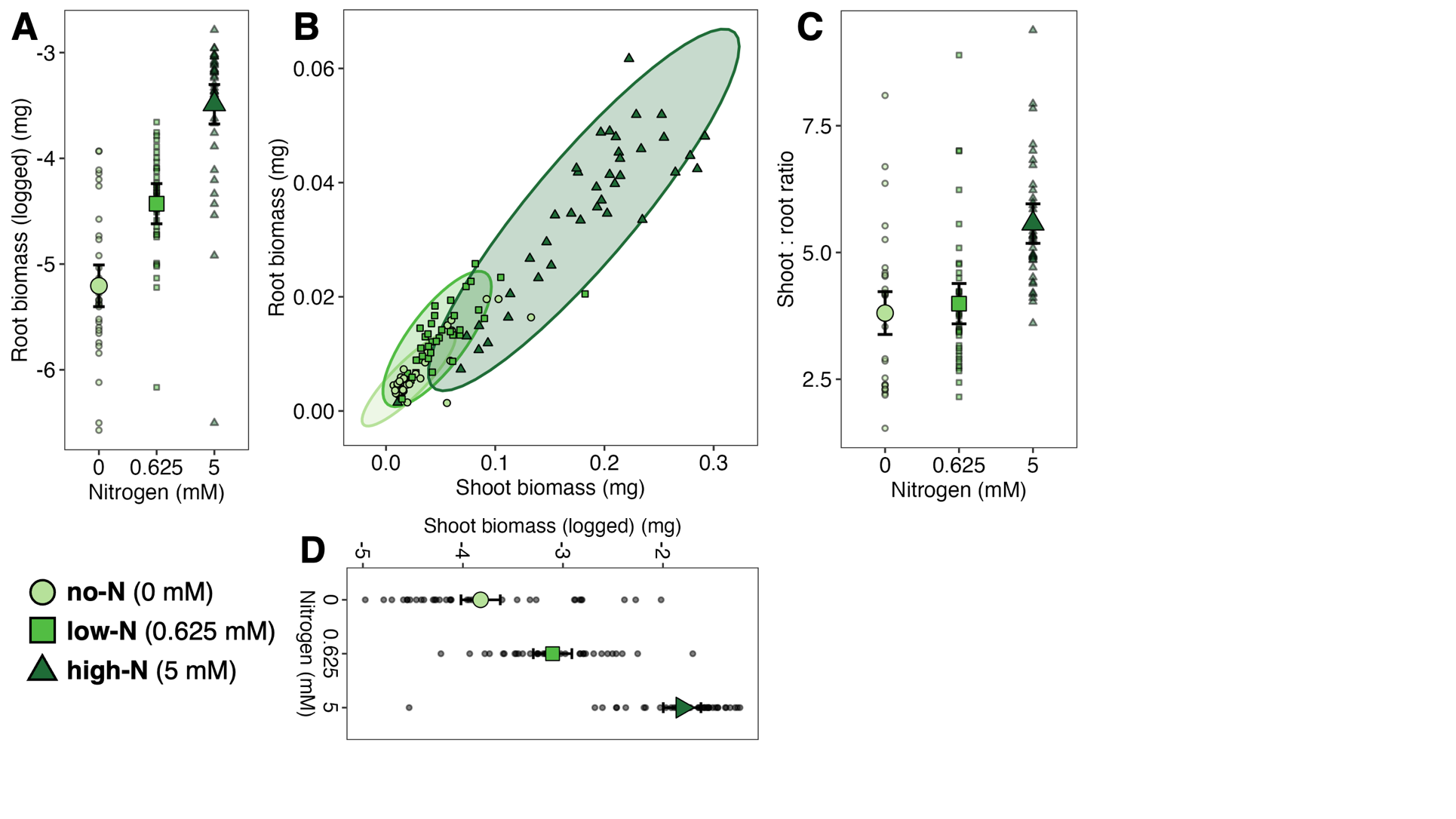

**Figure S3:** The effect of nitrogen treatment on logged root biomass, logged shoot biomass, and shoot:root ratios. Smaller transparent points are raw data, while the larger opaque points are the estimated marginal means. Error bars on the estimated marginal means show the 95% confidence interval.

**A.** Nitrogen treatment had a significant effect on logged root biomass (p < 0.001; Table S3A).

**B.** The relationship of root biomass to shoot biomass for each nitrogen treatment.

**C.** Nitrogen and block had a significant effect on shoot:root ratio. Rhizobia strain had a marginally significant effect on shoot:root ratio (p=0.0578; see Table S2B).

**D.** Nitrogen treatment had a significant effect on logged shoot biomass (p < 0.001; Table S3B).

**Table S3:** The effect of nitrogen treatment on shoots and roots.

|  | **X ^2^** | **Df** | **P** |
| --- | --- | --- | --- |
| A. **Effect of rhizobia strain and nitrogen on logged root biomass** | | | |
| **Nitrogen** | 153 | 2 | **<0.001** |
| Rhizobia | 3.90 | 4 | 0.420 |
| Block | 2.10 | 3 | 0.551 |
| Nitrogen:Rhizobia | 8.96 | 8 | 0.345 |
| B. **Effect of rhizobia strain and nitrogen on logged shoot biomass** | | | |
| **Nitrogen** | 224 | 2 | **<0.001** |
| Rhizobia | 7.00 | 4 | 0.136 |
| Block | 2.96 | 3 | 0.398 |
| Nitrogen:Rhizobia | 9.56 | 8 | 0.298 |

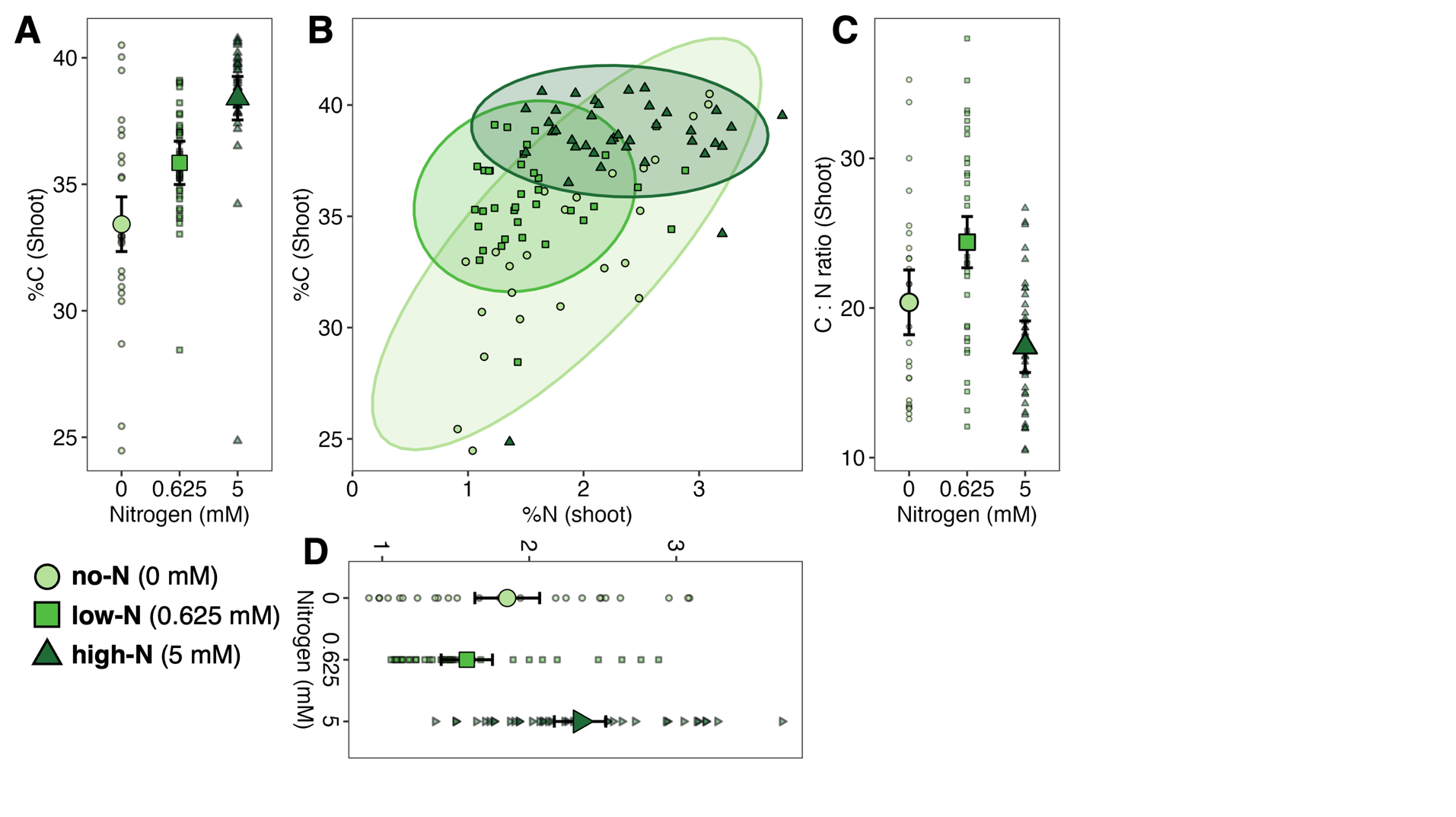

**Figure S4:** The effect of nitrogen treatment on shoot carbon and nitrogen and carbon:nitrogen use efficiency. Smaller transparent points are raw data, while the larger opaque points are the estimated marginal means. Error bars on the estimated marginal means show the 95% confidence interval.

**A.** Nitrogen treatment had a significant effect on shoot carbon concentration (p < 0.001; Table S4A).

**B.** The relationship of carbon to nitrogen concentration for each nitrogen treatment.

**C.** Nitrogen treatment had a significant effect of nitrogen on carbon:nitrogen ratio (p < 0.001; Table S4C). Block also had a significant effect on C:N ratio (p=0.0438).

**D.** Nitrogen treatment had a significant effect on on shoot nitrogen concentration (p < 0.001; Table S4B).

**Table S4:** The effect of nitrogen treatment on host carbon and nitrogen concentration.

|  | **X ^2^** | **Df** | **P** |
| --- | --- | --- | --- |
| A. **Effect of rhizobia strain and nitrogen on shoot carbon concentration** | | | |
| **Nitrogen** | 52.7 | 2 | **<0.001** |
| Rhizobia | 6.29 | 4 | 0.179 |
| Block | 3.44 | 3 | 0.328 |
| Nitrogen:Rhizobia | 8.93 | 8 | 0.349 |
| B. **Effect of rhizobia strain and nitrogen on shoot nitrogen concentration** | | | |
| **Nitrogen** | 39.1 | 2 | **<0.001** |
| Rhizobia | 3.86 | 4 | 0.425 |
| Block | 3.71 | 3 | 0.295 |
| Nitrogen:Rhizobia | 10.6 | 8 | 0.222 |
| C. **Effect of rhizobia strain and nitrogen on carbon:nitrogen ratio** | | | |
| **Nitrogen** | 32.9 | 2 | **<0.001** |
| Rhizobia | 3.36 | 4 | 0.500 |
| **Block** | 8.11 | 3 | **0.0438** |
| Nitrogen:Rhizobia | 10.4 | 8 | 0.236 |

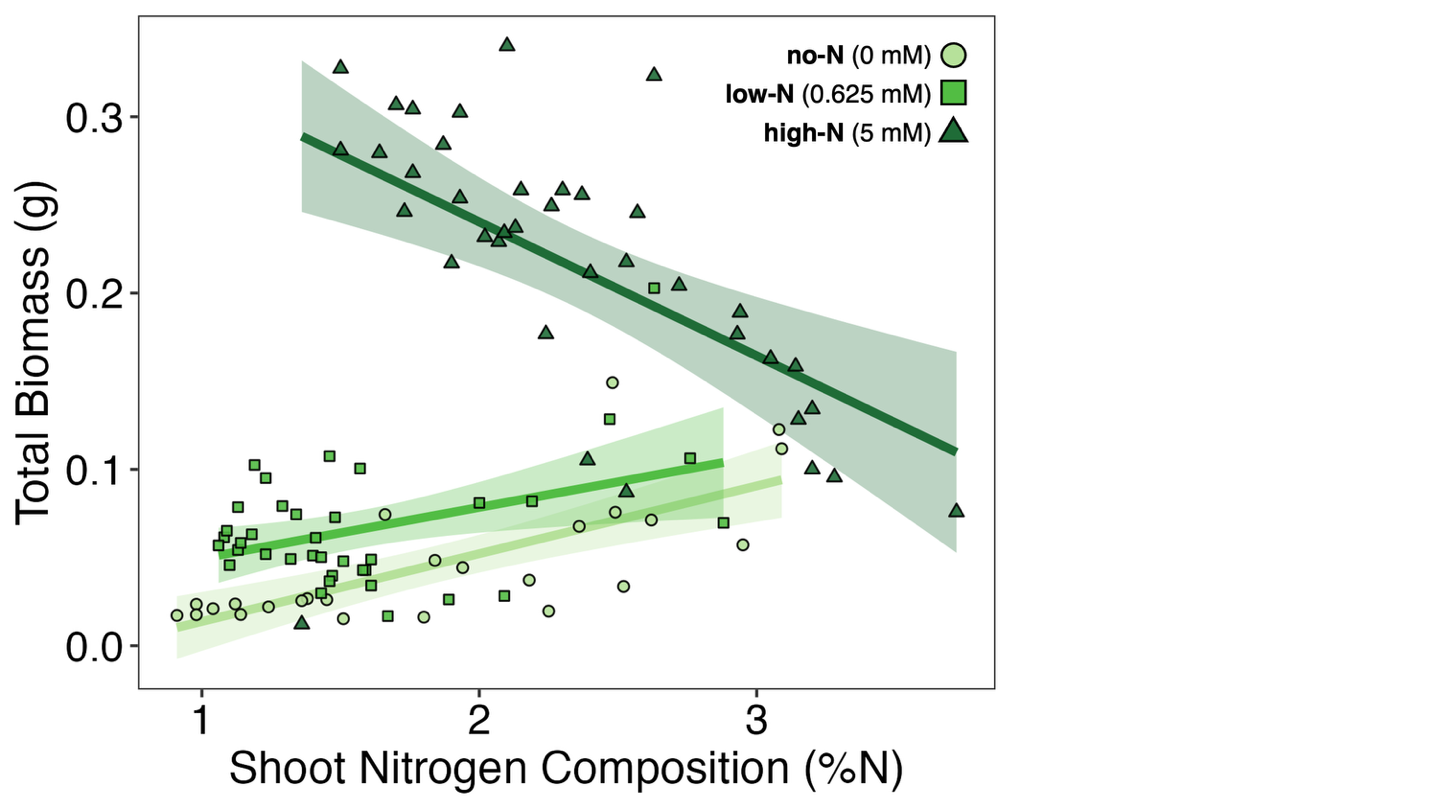

**Figure S5:** The relationship between total biomass and nitrogen concentration for each nitrogen treatment. For the high-N treatment, a negative slope suggests a tradeoff between total biomass and nitrogen concentration.

**Table S5:** The effect of rhizobia strain and nitrogen on total parasite load and size-adjusted parasite load (intensity of infection). Paired with Figure 1 in main text.

|  | **A: The effect of rhizobia strain and nitrogen on total parasite load** | | | **B: The effect of rhizobia strain and nitrogen on size-adjusted parasite load (intensity of infection)** | | |
| --- | --- | --- | --- | --- | --- | --- |
|  | **X ^2^** | **Df** | **P** | **𝝌^2^** | **Df** | **P** |
| Rhizobia | 2.57 | 4 | 0.631 | 0.809 | 4 | 0.937 |
| **Nitrogen** | 40.0 | 2 | **<0.001** | 28.2 | 2 | **<0.001** |
| Block | 4.55 | 3 | 0.208 | 2.30 | 3 | 0.513 |
| Rhizobia:Nitrogen | 9.86 | 8 | 0.275 | 7.98 | 8 | 0.435 |
| **Root biomass  (z-score)** | NA | NA | NA | 26.2 | 1 | **<0.001** |

**Table S6:** Post-hoc tests of differences between nitrogen treatments when testing for the effect of nitrogen and rhizobia inoculation on total parasite load. Paired with Figure 1 in main text.

|  | **A: The effect of nitrogen and nodulation on total parasite load** | | | **The effect of nitrogen and nodulation on size-adjusted parasite load (intensity of infection)** | | |
| --- | --- | --- | --- | --- | --- | --- |
|  | **Ratio** | **SE** | **P** | **Ratio** | **SE** | **P** |
| No-N and Low-N  (0 - 0.625 mM) | 0.324 | 0.0984 | **<0.001** | 0.398 | 0.116 | **0.0043** |
| No-N and High-N  (0 - 5 mM) | 0.270 | 0.0900 | **<0.001** | 0.996 | 0.391 | 1.000 |
| Low-N and High-N  (0.625 - 5 mM) | 0.834 | 0.252 | 0.820 | 2.51 | 0.846 | **0.0178** |

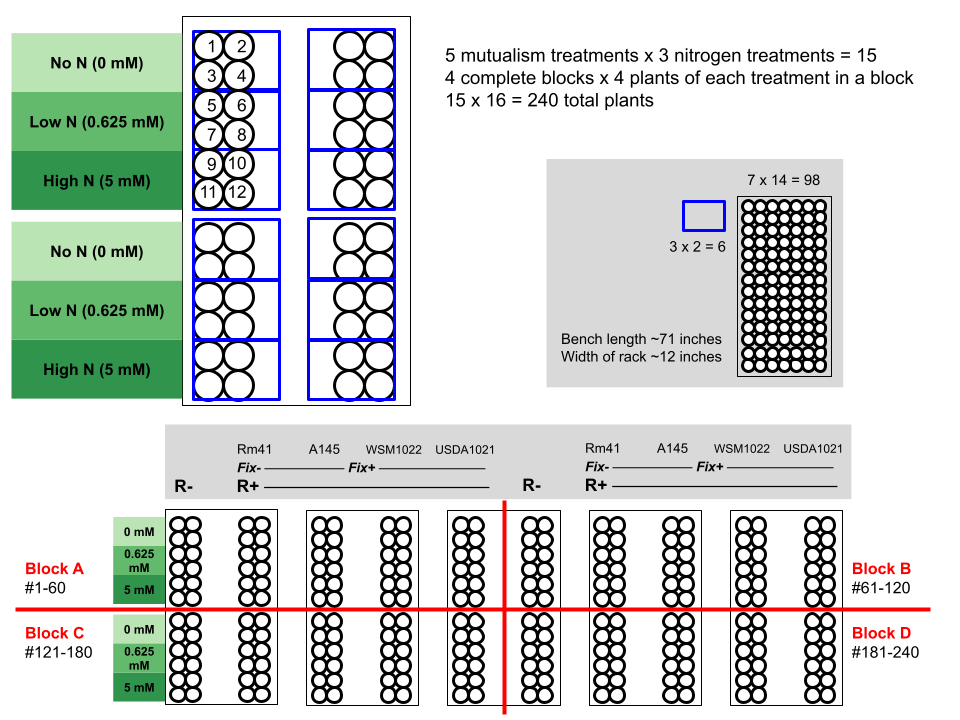

**Figure S6:** The full blocking scheme for the greenhouse experiment included five Cone-tainer^TM^ racks split into four complete blocks. Each rack contained two rhizobia treatments and all three nitrogen fertilizer treatments; each block contained all five rhizobia treatments and all three nitrogen treatments. Plant IDs were numbered left to right and top to bottom within each spatially separated group of 12 plants.

**Table S7:** Post-hoc tests of differences between nitrogen treatments on parasite load after adjusting for the effect of N-fixing and non-N-fixing nodules, block, and the interaction between nitrogen treatments and each nodule type. Paired with Figure 2 in main text.

|  | **Ratio** | **SE** | **P** |
| --- | --- | --- | --- |
| **No-N and Low-N**  **(0 - 0.625 mM)** | 0.249 | 0.0614 | **<0.001** |
| **No-N and High-N**  **(0 - 5 mM)** | 0.185 | 0.0470 | **<0.001** |
| Low-N and High-N  (0.625 - 5 mM) | 0.741 | 0.1270 | 0.1890 |

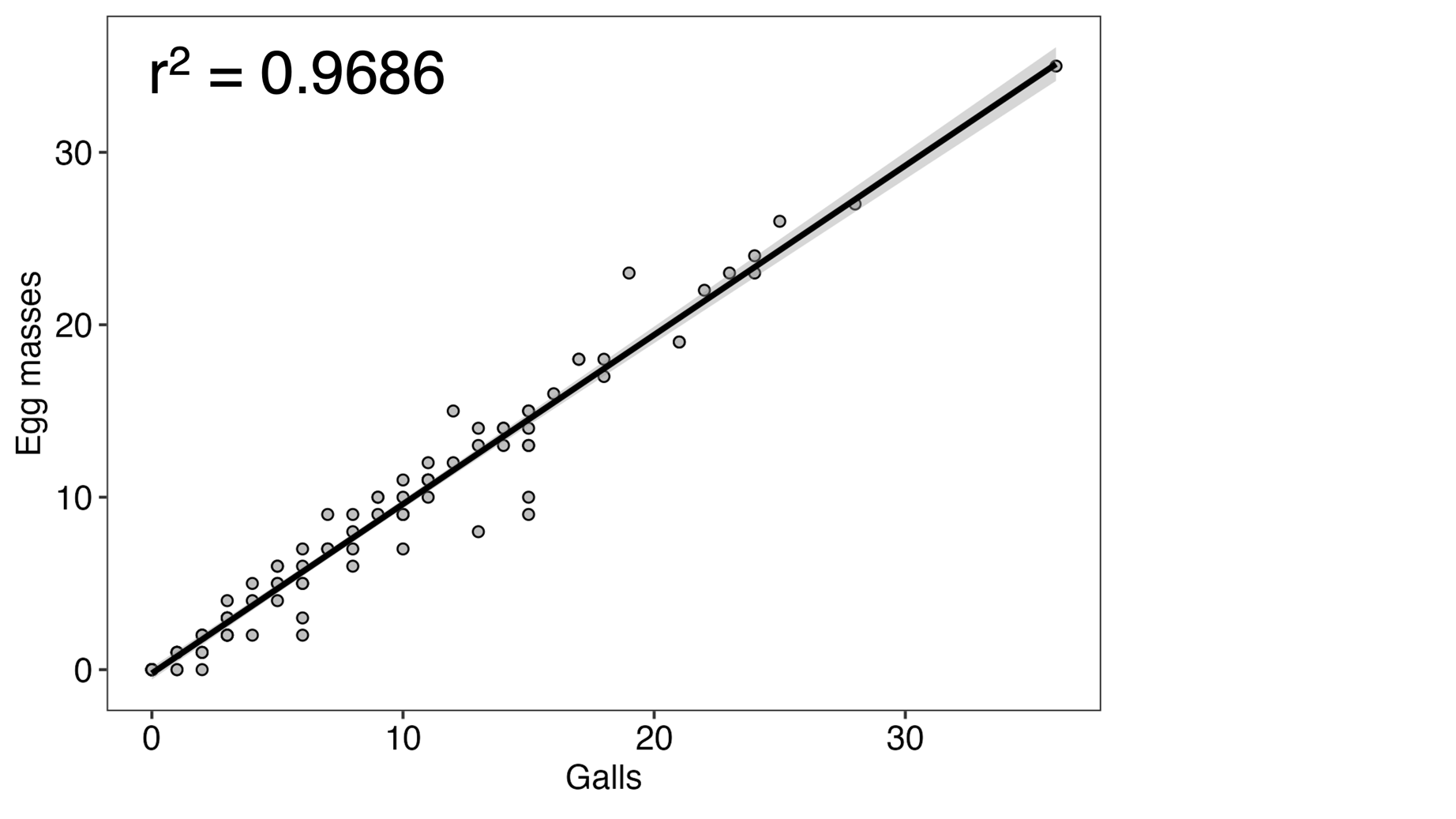

**Figure S7:** The relationship between galls and egg masses. R-squared value was derived using the summary function (r.squared) on a simple linear regression of galls on egg masses.

**Table S8:** Post-hoc tests of differences between nitrogen treatments on parasite load after adjusting for the effect of nitrogen, rhizobia, root biomass, shoot nitrogen concentration, and shoot carbon concentration. Paired with Figure 3 in main text.

|  | **Ratio** | **SE** | **P** |
| --- | --- | --- | --- |
| **No-N and Low-N**  **(0 - 0.625 mM)** | -1.31 | 0.263 | **<0.001** |
| **No-N and High-N**  **(0 - 5 mM)** | -1.39 | 0.268 | **<0.001** |
| Low-N and High-N  (0.625 - 5 mM) | -0.0729 | 0.154 | 0.884 |

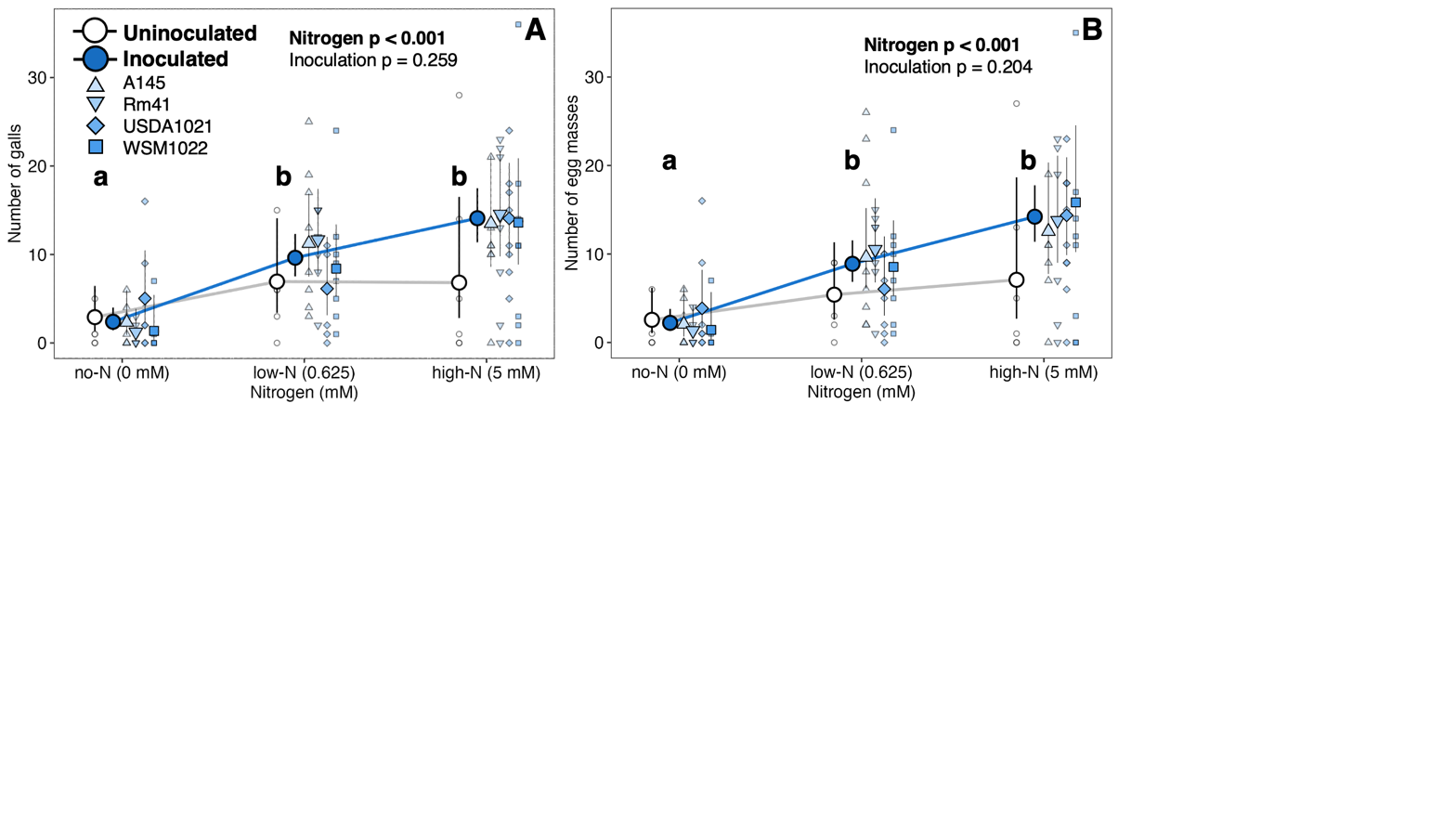

**Figure S8:** A comparison of two measures of root-knot nematode infection by nitrogen fertilizer and rhizobia inoculation treatments. Responses are visually similar.

**A.** We counted the number of gall root structures as a measure of parasite load.

**B.** We counted the number of egg masses: each reproductive female generally lays one egg mass.

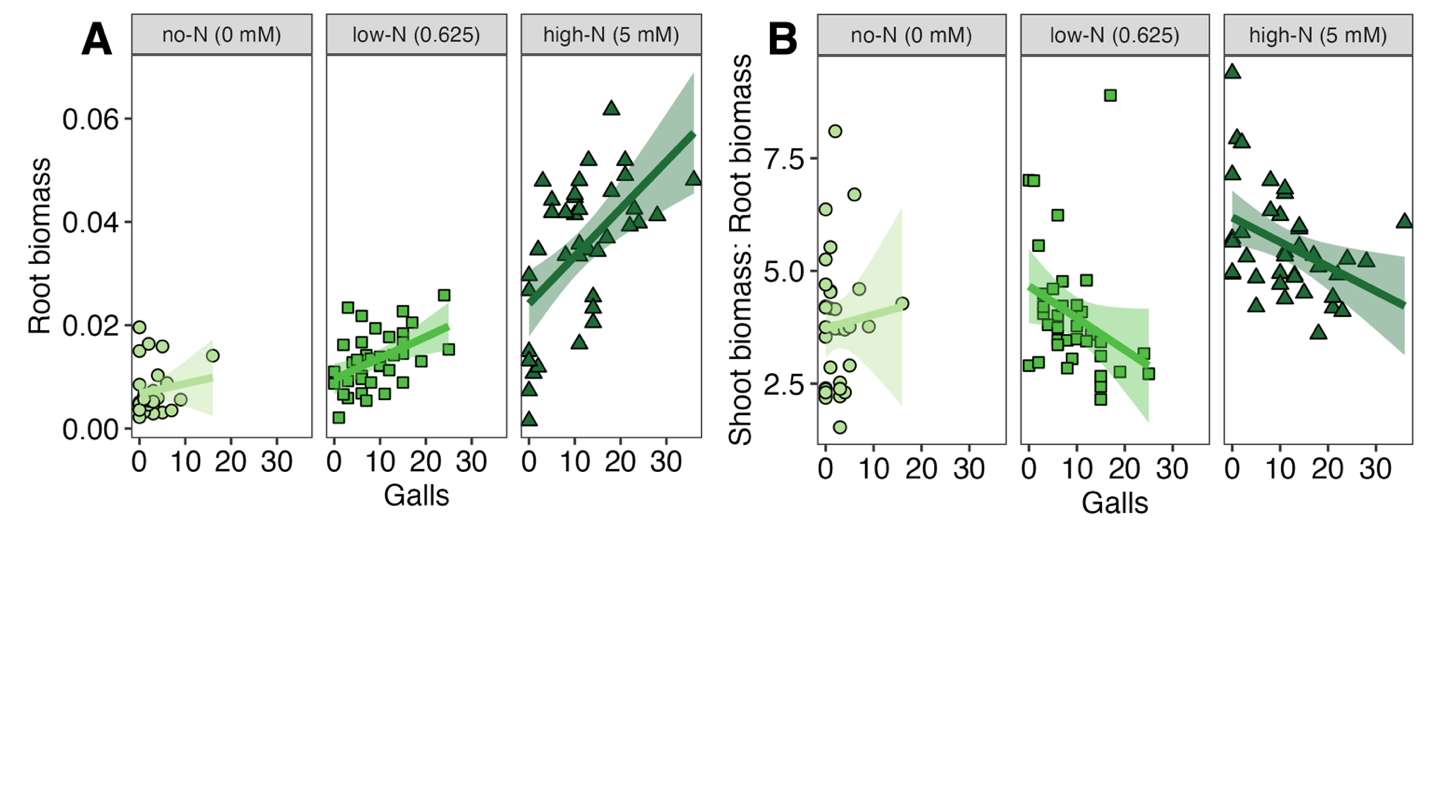

**Figure S9:** The relationship between galls and biomass allocation.

A. Root biomass and gall count, faceted across nitrogen treatments.

B. Shoot:root ratios and gall count, faceted across nitrogen treatments.

**Table S9:** Recipes to make fertilizer solutions (adapted from Batstone *et al.* 2017, originally based on table 2 of Moreau et al., 2008) at various N-concentrations. Bolded compounds differ in concentration across fertilizer types. * Iron versenate (Fe-EDTA) was made by mixing FeSO4 (5.6 g/l) and Na2-EDTA (7.4 g/l) at 50 °C. NA refers to not applicable (no volume added).

|  |  |  | **1L 0 mM-N fertilizer (no-N)** | | **1L 0.625 mM-N fertilizer (low-N)** | | **1L 5 mM-N fertilizer (high-N)** | |
| --- | --- | --- | --- | --- | --- | --- | --- | --- |
|  | **Compound** | **Final concentration (mol/L)** | **mL** | **mmol/L (mM)** | **mL** | **mmol/L (mM)** | **mL** | **mmol/L (mM)** |
| Macronutrients | MgSO4 • 7 H2O | 0.500 | 4.000 | 2.0000 | 4.000 | 2.0000 | 4.000 | 2.0000 |
|  | **CaCl2 • 2 H2O** | 1.000 | **5.000** | **5.0000** | **4.530** | **4.5300** | **3.125** | **3.1250** |
|  | Fe-EDTA* | 0.020 | 2.500 | 0.0500 | 2.500 | 0.0500 | 2.500 | 0.0500 |
| Micronutrients | MnSO4 | 0.006 | 0.100 | 0.0006 | 0.100 | 0.0006 | 0.100 | 0.0006 |
|  | CuSO4 | 0.006 | 0.100 | 0.0006 | 0.100 | 0.0006 | 0.100 | 0.0006 |
|  | ZnSO4 (0.1M) | 0.006 | 0.100 | 0.0006 | 0.100 | 0.0006 | 0.100 | 0.0006 |
|  | H3BO3 | 0.016 | 0.100 | 0.0016 | 0.100 | 0.0016 | 0.100 | 0.0016 |
|  | Na2MoO4 | 0.005 | 0.100 | 0.0005 | 0.100 | 0.0005 | 0.100 | 0.0005 |
| Nitrogen | **KNO3** | 0.625 | NA | NA | **0.250** | **0.1560** | **1.000** | **0.6250** |
|  | **Ca(NO3)2 • 4 H2O** | 0.938 | NA | NA | **0.500** | **0.4690** | **2.000** | **1.8750** |
|  | **NaNO3** | 2.500 | NA | NA | NA | NA | **1.000** | **2.5000** |
| Fertilizer | K2HPO4 | 1.200 | 2.000 | 2.4000 | 2.000 | 2.4000 | 2.000 | 2.4000 |
|  | **K2SO4** | 0.430 | **2.197** | **0.9456** | **1.953** | **0.8406** | **1.802** | **0.7756** |
|  | NaCl | 0.200 | 1.000 | 0.2000 | 1.000 | 0.2000 | 1.000 | 0.2000 |
| Water | H2O |  | 982.803 |  | 982.767 |  | 981.073 |  |

**Table S10**: The effect of nitrogen and rhizobia inoculation (A) and strain (B) on two measures of parasite load (galls and egg masses). Responses are statistically similar; galls and egg masses are highly correlated (r^2^=0.9686).

|  | **A. The effect of nitrogen and rhizobia inoculation on parasite load** | | | | | |
| --- | --- | --- | --- | --- | --- | --- |
|  | **Galls as response variable (measure of parasite load)** | | | **Egg masses as response variable (measure of parasite load)** | | |
|  | **𝝌^2^** | **Df** | **P** | **𝝌^2^** | **Df** | **P** |
| Inoculation | 1.2732 | 1 | 0.259 | 1.6141 | 1 | 0.2039 |
| **Nitrogen** | 18.855 | 1 | **<0.0001** | 17.257 | 1 | 0.000179 |
| Block | 5.5127 | **2** | 0.138 | 4.5852 | **2** | 0.205 |
| Inoculation:Nitrogen | 1.9491 | 3 | 0.377 | 1.5179 | 3 | 0.468 |
|  | **B. The effect of nitrogen and rhizobia strain on parasite load** | | | | | |
|  | **Galls as response variable (measure of parasite load)** | | | **Egg masses as response variable (measure of parasite load)** | | |
|  | **𝝌^2^** | **Df** | **P** | **𝝌^2^** | **Df** | **P** |
| Rhizobia | 1.7502 | 4 | 0.6308 | 1.7502 | 4 | 0.7816 |
| **Nitrogen** | **39.5134** | **2** | **<0.0001** | **39.513** | **2** | **<0.0001** |
| Block | 4.5534 | 3 | 0.2079 | 4.5534 | 3 | 0.2076 |
| Rhizobia:Nitrogen | 6.7424 | 8 | 0.2747 | 6.7424 | 8 | 0.5647 |

**Table S11**:  **A.** The effect of nitrogen and inoculation treatments on parasite load, comparing between 20 uninoculated (R-) plants and 30 plants inoculated with a single focal strain (USDA1021).  **B.** Post-hoc tests of differences between nitrogen treatments on parasite load for the effect of nitrogen, rhizobia, root biomass, shoot nitrogen concentration, and shoot carbon concentration between 20 uninoculated (R-) plants and 30 plants inoculated with a single focal strain (USDA1021).

|  | **A. The effect of inoculation (R- vs USDA1021) and nitrogen on total parasite load** | | |
| --- | --- | --- | --- |
|  | **Chisq** | **Df** | **Pr(>Chisq)** |
| Inoculation | 0.1158 | 1 | 0.7336 |
| **Nitrogen** | 19.8856 | 2 | **<0.001** |
| Block | 6.5480 | 3 | 0.0879 |
| Inoculation:Nitrogen | 1.1352 | 2 | 0.56689 |
|  | **B. Post-hoc differences between uninoculated R- plants and USDA1021 as focal strain on parasite load** | | |
|  | **Ratio** | **SE** | **P** |
| At no-N (0 mM) | 0.643 | 0.326 | 0.3830 |
| At low-N (0.625 mM) | 1.316 | 0.574 | 0.5296 |
| At high-N (5 mM) | 0.914 | 0.320 | 0.7976 |
